# Uncovering Ant Diversity Across Forest Successional Stages in the Yangambi Biosphere Reserve (DRC): Insights from Winkler and Pitfall Trap Sampling

**DOI:** 10.64898/2025.12.04.692217

**Authors:** Hervé Ambakina, Kiko Gomez, Jean-Claude Monzenga, Daniel Sánchez-García, Wouter Dekoninck

## Abstract

Litter ant communities are an important component of biodiversity in tropical regions. They are currently used in several ecosystem management programmes to assess forest health. The aim of this study was to uncover the ant diversity across forest successional stages (fallow, secondary forest and primary forest) in the Yangambi Biosphere Reserve, The Democratic Republic of the Congo. These three forest habitats were sampled in six localities with pitfall traps and Winkler extractions. The discovered ant communities, the species typical of each habitat and the effectiveness of both sampling methods are discussed. The results showed that 190 species belonging to 50 genera and 8 sub-families of Formicidae were sampled in the Yangambi region. Ant diversity was lowest in fallow land, higher in secondary forests, and highest in primary forests, reflecting a clear pattern along the successional gradient. A relatively small proportion of ant species were shared among all three habitats, while each habitat also supported a distinct set of species. Primary forests hosted the highest number of exclusive species, followed by secondary forests and fallow land. Winkler extractors captured substantially more ant species than pitfall traps, with nearly 50% greater species richness observed. However, a significant portion of the ant fauna in Yangambi likely remains undocumented, and additional sampling methods could help provide a more complete picture of its biodiversity.

## INTRODUCTION

Tropical forests are of exceptional value for biological conservation (WWF 2007; Fao 2010; Marsh et al. 2025). They are home to more than half of the world’s living species, including plants, animals, insects and micro-organisms and are vital to the preservation of biological diversity (Pillay et al. 2021; Boyle et al. 2025; Li 2025). These forests are currently being destroyed at an estimated rate of around 10 million ha per year (Laso Bayas et al. 2022). Several researchers around the world have already shown that the loss or disappearance of species worldwide is mainly due to the destruction of wooded vegetation cover (Giam 2017) and may drive the extinction of approximately a quarter of global species within the next few decades. (Tilman et al. 2017; Strona and Bradshaw 2022).

Forests in the Afrotropics face the same threats. For example, in the Democratic Republic of the Congo (DRC), forests face not only intense pressure from a growing population but also pervasive industrial threats. The DRC is experiencing an accelerating rate of deforestation, especially in the Yangambi Biosphere Reserve (Mayaux et al. 2003), largely due to regional population growth (WWF 2007). Consequently, several plant formations are emerging in the Yangambi Biosphere Reserve, a process believed to promote biological erosion (Alongo 2007, 2013). Primary forest is continuously being transformed to fallow land which later -after reconversion and restauration-might lead to parcels with secondary forest. Estimating the species richness of such a given area like transformed or primary forests is challenging, as existing inventory methods often fail to capture a representative sample of the total diversity. There is indeed a need for well-structured sampling protocols to properly estimate these species richness. For ants in tropical forest the ALL (Ants of the Leaf Litter) protocol was developed and often used (Agosti and Alonso 2000). This protocol is a standardized method used to sample ant diversity, especially in leaf litter habitats. It typically involves: leaf litter collection (collecting litter from a defined area (usually 1 m²)) and consequently sifting (using a mesh sieve to remove large debris and concentrate the fine litter containing ants and other small invertebrates) and extraction (placing the sifted litter into a Berlese or Winkler extractor, which uses heat and/or drying to drive ants down into a collecting container (usually with alcohol for preservation). While ALL focuses heavily on leaf litter sampling using Winkler extractors, it often also incorporates multiple complementary methods, including Pitfall traps (for surface-active ants), hand collecting (for visible foragers or nests) and baiting (to attract certain ant species).

In this study, we investigate leaf litter ant communities in different habitats of the Yangambi Biosphere Reserve using Winkler extraction and pitfall traps, which are both evaluated and compared to determine their effectiveness in sampling ant diversity. Additionally, we seek to determine whether the species richness of primary forest is truly comparable to that of secondary forests and fallow areas. Moreover, we also want to understand whether the destruction of forest habitats contributes to the loss of ant fauna of the leaf litter and to analyse whether the reconstitution of the forest also contributes to the conservation of biological diversity.

## MATERIAL AND METHODS

### Study area

The Yangambi forest region is located at 0°45’ N, 24°29’ E, and an altitude of 500 meters above sea level (Bernard 1945; De Leenheer et al. 1952) covering approximately 6.297 km2 (Drachossoff et al. 1991) in DRC. Current climate is hot, humid continental equatorial (Bernard 1945) and of the Af de Köppen type (for more explanation see Beck et al. 2018). The region has an average annual temperature of 24.6°C (Vandenput 1981), an average annual rainfall of 1875 mm, and characteristic ferralitic soils (Kombele 2004). Plant diversity is remarkably high, ranging from the pioneer stage including forest regrowth, secondary and disturbed forests to climax forests including semi-deciduous forests, *Gilbertiodendron dewevrei* rainforests, *Brachystegia laurentii* climatic forests, as well as riparian and swamp forests. In the secondary rain forest *Pycnanthus angolensis* and *Fagara macrophylla* are the dominating plant species (www.yangambi.org).

### Sampling

The study was carried out from November 2023 to February 2024 combining Winkler extractions and pitfall traps. Six different locations in the Yangambi Biosphere Reserve were selected: Bangala, Gazi, Ifa, Likango, Lumumba and Musa (Figure 1). To assess ant diversity across different forest successional stages, three habitat types—fallow, secondary forest, and primary forest—were sampled at each designated location, considering them as study plots. In each plot, sampling effort consisted of four independent transects. Each transect incorporated five pitfall traps spaced 10 m apart and five 1 m^2^ litter samples collected at corresponding 10 m intervals along the same transects.

**Figure 1.**
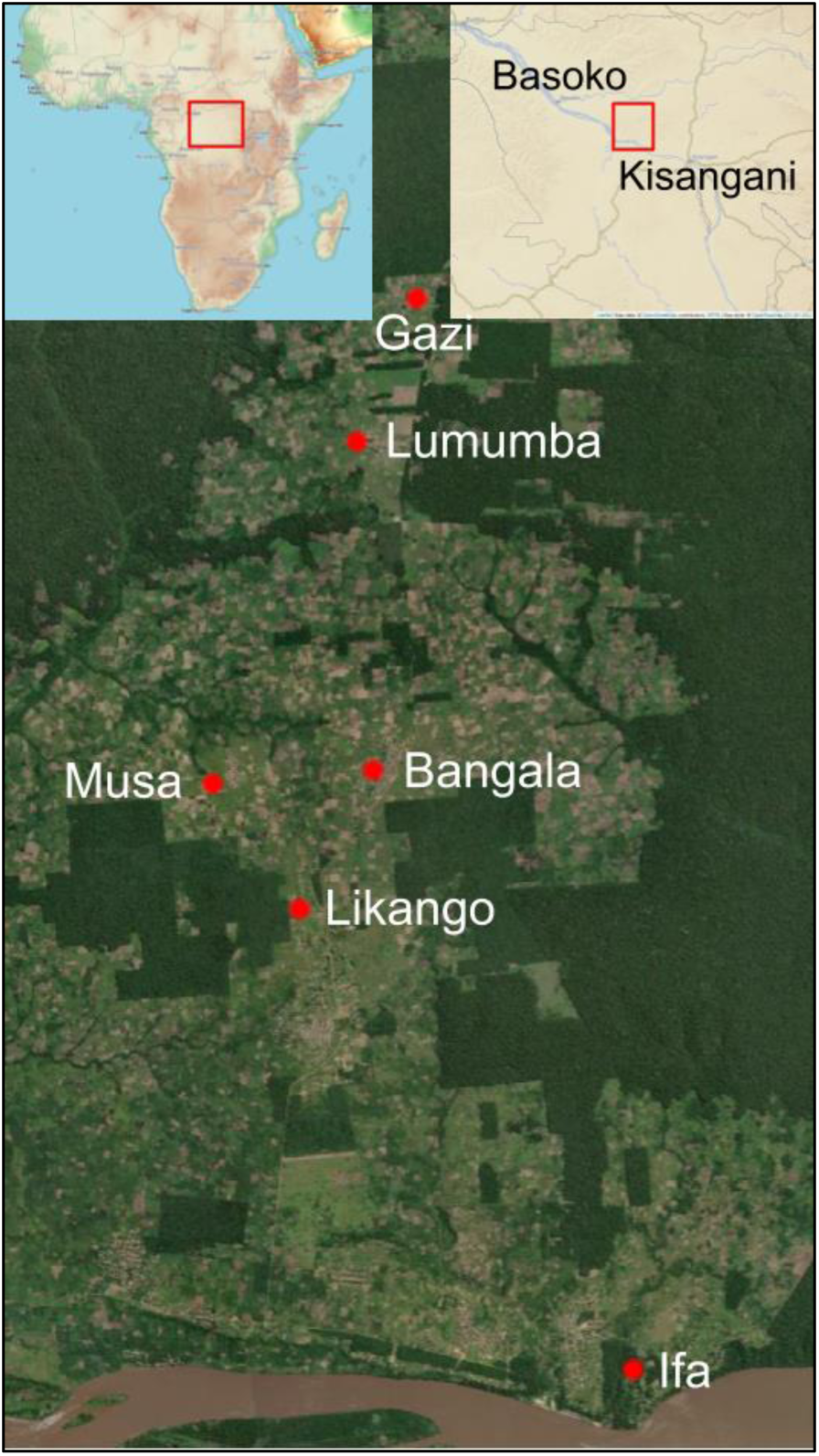
Sampled locations in the Yangambi Forest Reserve

Litter was sieved with a 1 cm mesh size sieve in the field. This technique reduces litter and concentrates the fauna (Agosti and Alonso 2000). Leaf litter was transferred into small, perforated bags and suspended in the mini-Winkler bags for 48 hours to extract the fauna. During this drying process, the fauna inside the Winkler starts to migrate downwards and falls into a pot containing 70% ethanol. Pitfall traps were placed in the same spots along the transact. The traps consisted of pots of 12 cm high and 8 cm in diameter, containing soapy water and salt filled for three-fourths. Each pitfall was protected by a plant leaf to prevent rainwater and all sorts of plant debris that could damage the capture. Samples were stored in 70% ethanol.

### Identification and storage

Ants are one of the most indispensable groups of insects in tropical regions and consequently also in forests. The ant community alone accounts for more than a third of the total insect biomass (Hölldobler and Wilson 1990) and its total number being estimated in 20 quadrillion (Schultheiss et al. 2022). Such key ant populations regulate the overpopulation of other living organisms in ecosystems (Bronstein et al. 2006; Dejean et al. 2007). Ants contribute to the trophic chain and stimulates the decomposition of organic matter on the soil surface. This community is one of the robust bioindicators frequently used to assess biotic and ecosystem responses to environmental change (Agosti et al. 2000; Andersen et al. 2002; Andersen and Majer 2004; Underwood and Fisher 2006; Lapolla et al. 2006; Rohr et al. 2006; Majer et al. 2007).

All ant samples were sent to the Royal Belgian Institute of Natural Sciences (RBINS) for identification. Voucher samples are stored in RBINS and the private collection of the second author; Kiko Gomez Ant Collection (KGAC). Voucher specimens will also be stored in DRC local collections to build a local reliable identification source for future myrmecological studies.

Identification to genus level was conducted with Fisher and Bolton, 2016. For the identification at species level we used the following keys: *Aenictus* (Gomez 2022), *Anochetus* (Brown 1978), *Baracidris* (Fernandez 2003), *Bothroponera* (Joma & Mackay 2015, 2020), *Calyptomyrmex* (Bolton 1981a), *Carebara* (former *Paedalgus* (Bolton and Belshaw 1993), group *polita* (Fischer et al. 2014), *Cataulacus* (Bolton 1982), *Dicroaspis* (Bolton 1981a), *Discothyrea* (Hita Garcia et al. 2019), *Eburopone* (Yamada et al. 2023), *Hypoponera* (Bolton and Fisher 2011), *Leptogenys* (Bolton 1975), *Meranoplus* (Bolton 1981), *Microdaceton* (Bolton 2000), *Monomorium* (Bolton 1987), *Nesomyrmex* (Hita Garcia et al. 2017), *Nylanderia* (Lapolla et al. 2011), *Odontomachus* (Brown 1976), *Paraparatrechina* (Lapolla and Fisher 2014), *Paratrechina* (Lapolla et al. 2013), *Pheidole* group *pulchella* (Gomez et al. 2022), *Phrynoponera* (Bolton and Fisher 2008), *Platythyrea* (Brown 1975), *Polyrhachis* (Rigato 2016), *Pristomyrmex* (Wang 2003), *Strumigenys* (Bolton 2000), *Syllophopsis* (Bolton 1987), *Technomyrmex* (Bolton 2007), *Tetramorium* (Bolton 1976, 1980, Hita Garcia et al. 2010; Hita Garcia and Fisher 2014) and *Tetraponera* (Ward 2022).

Non-revised genera were identified by comparing these specimens with type specimens available at antweb.org or specimens stored in RBINS and KGAC. This was done for specimens belonging to the following genera: *Anoplolepis*, *Camponotus*, *Carebara* (remainder of groups), *Crematogaster*, *Dorylus*, *Fisheropone*, *Lepisiota*, *Lioponera*, *Megaponera*, *Mesoponera*, *Myrmicaria*, *Oecophylla*, *Paltothyreus*, *Parasyscia*, *Pheidole* (remainder of groups), *Plagiolepis*, *Prionopelta*, *Probolomyrmex*, *Stigmatomma* and *Tapinoma*.

When no satisfactory match was found, the morphospecies were given a code for databasing purposes and are listed here as sp01, kgac-drc01, etc.

### Biodiversity indices and statistical analyses

Ant communities were analyzed based on species abundance, which was calculated as occurrence per transect, with a maximum of four occurrences per study plot (one per transect).

Species richness, Shannon diversity, and Simpson diversity indices were calculated using specnumber() and diversity() functions from the vegan package (Oksanen et al. 2025).

Differences in these indices among habitat types and sampling methods were tested using the Kruskal-Wallis test implemented in the rstatix package (Kassambara 2023), with pairwise comparisons conducted using Dunn’s test (Dinno 2024) with Holm’s and Benjamini-Hochberg’s adjustment.

Differences in ant community composition among habitat types and sampling methods were assessed based on Bray-Curtis dissimilarity using permutational multivariate analysis of variance (PERMANOVA) with 999 permutations implemented in the vegan package (Oksanen et al. 2025). Pairwise comparisons were conducted using the pairwiseAdonis package (Martinez Arbizu 2017) with Holm’s and Benjamini-Hochberg’s adjustment. Non-metric multidimensional scaling (NMDS) was performed using the metaMDS() function (Oksanen et al. 2025) based on the same Bray-Curtis dissimilarity distances to visualize differences in community composition. Homogeneity of dispersion (beta diversity) among habitats or sampling methods was assessed using the betadisper() function (Oksanen et al. 2025), using ANOVA (R Core Team 2024) to test for significant differences in dispersion, and performing pairwise comparisons using Tukey’s HSD test (R Core Team 2024).

Species accumulation curves were generated using the specaccum() function (Oksanen et al. 2025) with a random model generation approach and 999 permutations. Asymptotic models were fitted using fitspecaccum() (Oksanen et al. 2025) to estimate maximum species richness from the mean asymptote value.

Outlier fitted models were removed based on 3 standard deviations from the mean of the asymptotic value. We predicted the estimated species richness from the fitted model at our maximum sampling effort. We also estimated the sampling effort required to achieve 95 % and 99 % of maximum expected species richness, based on the fitted asymptotic models.

The log rate constant (lrc) was estimated and normalized by dividing the log rate constant between the maximum expected species richness of each rarefaction curve to be able to compare the rate of species accumulation among habitats and sampling methods. All statistical analyses were performed using R (<u>R Core Team 2024</u>).

## RESULTS

### General results on ant diversity and communities

A total of 190 ant species, representing 50 genera and 8 subfamilies, were collected using the two sampling methods (Table 1). Diversity indices of litter ant communities in the Yangambi region vary across habitat types (Kruskal-Wallis test | Specific richness (S): d.f. = 2, χ^2^ = 12.7, p = 0.002; Shannon index (H’): d.f. = 2, χ^2^ = 12.78, p = 0.002; Simpson index: d.f. = 2, χ^2^ = 12.54, p = 0.002). For all indices primary and secondary forests showed higher values compared to fallow communities (Figure 2).

**Figure 2.**
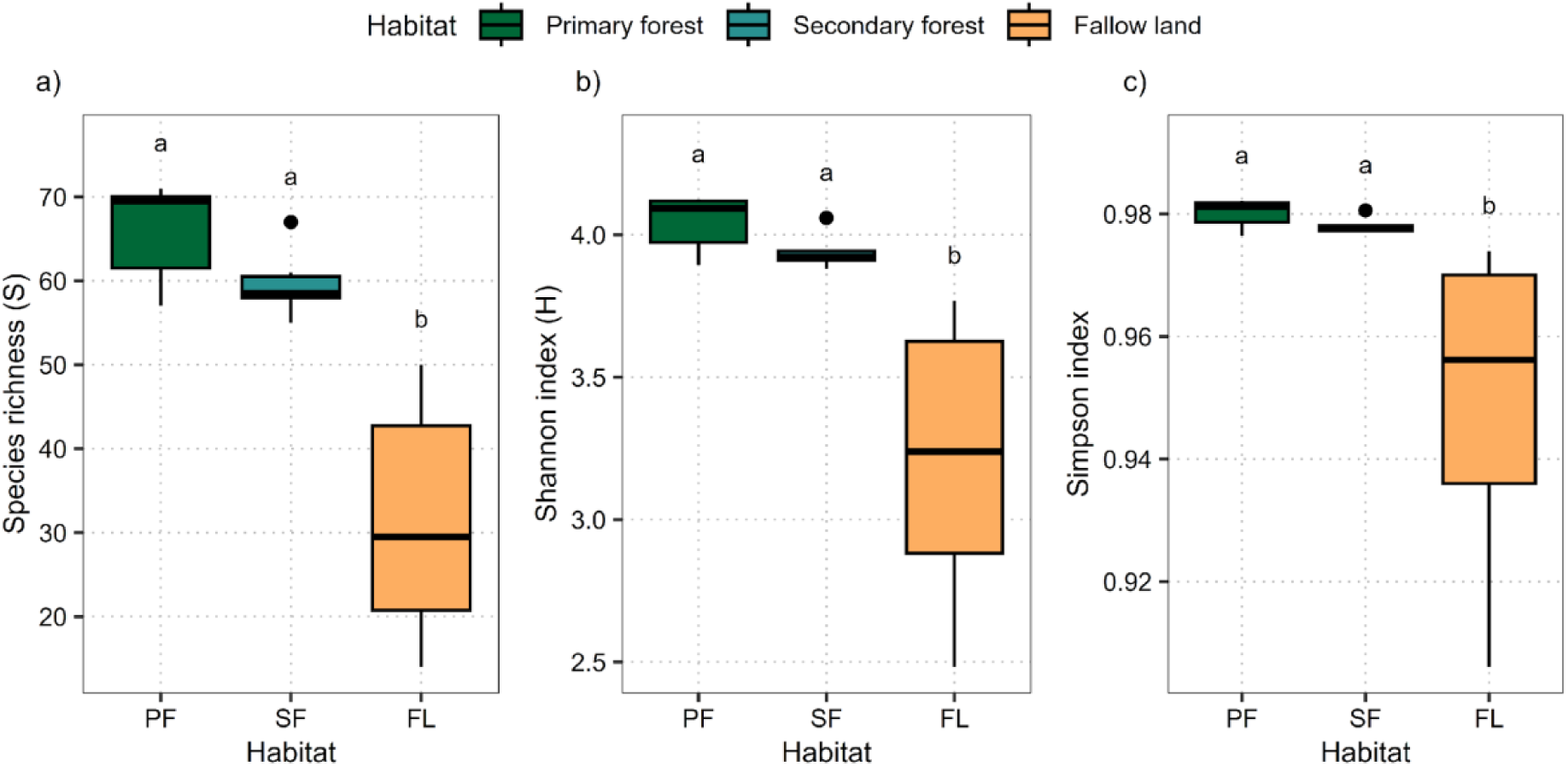
Litter ant community biodiversity indices in the different habitats Pairwise Dunn’s test results for the comparison of the effect of habitat type on diversity indices can be found in Table S1.

Litter ant communities differed in composition across the three habitats, with each forest type supporting its own set of unique species. Primary forests harboured the highest number of unique species, while fallow lands had the fewest. Although some species were shared among all habitats (60 common species), most overlap occurred between primary and secondary forests, with minimal shared species between fallow land and the other two habitats (Figure 3).

**Figure 3.**
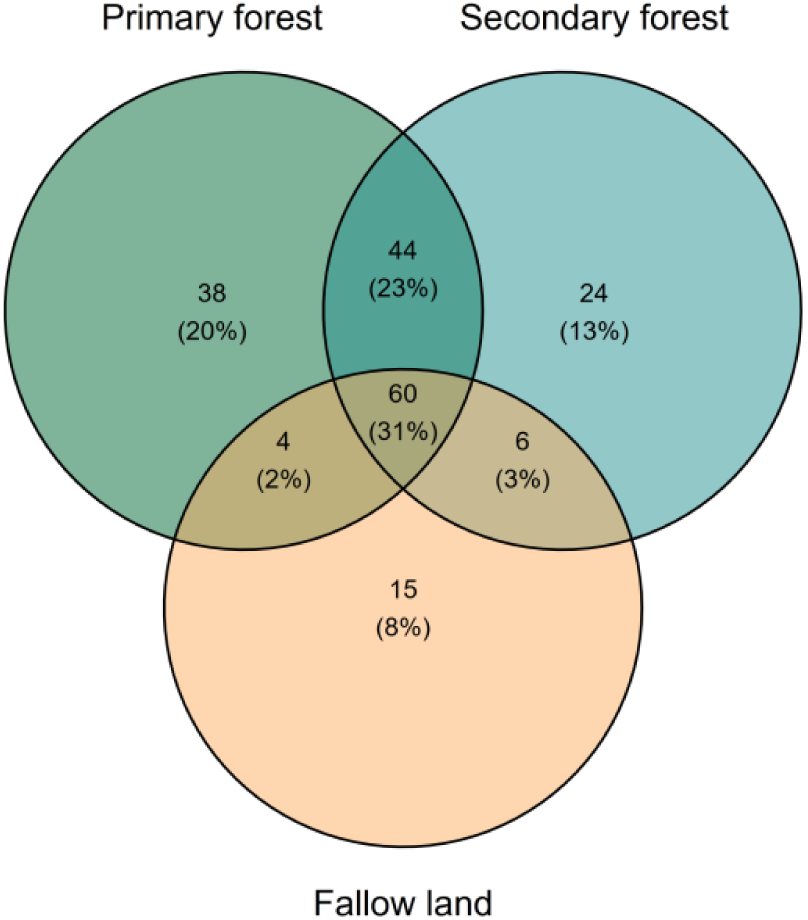
Venn diagram showing the number and percentage of species per habitat and habitat combination.

Ant communities differed significantly among habitats (PERMANOVA: d.f. = 2, F = 6.33, p < 0.001; Figure 4a). While primary and secondary forest did not show differences between their communities, fallow land differed significantly from both of them (Table S2). Additionally, fallow communities showed a higher variability between sites, compared to primary and secondary forest (Figure 4b; Table S3).

**Figure 4:**
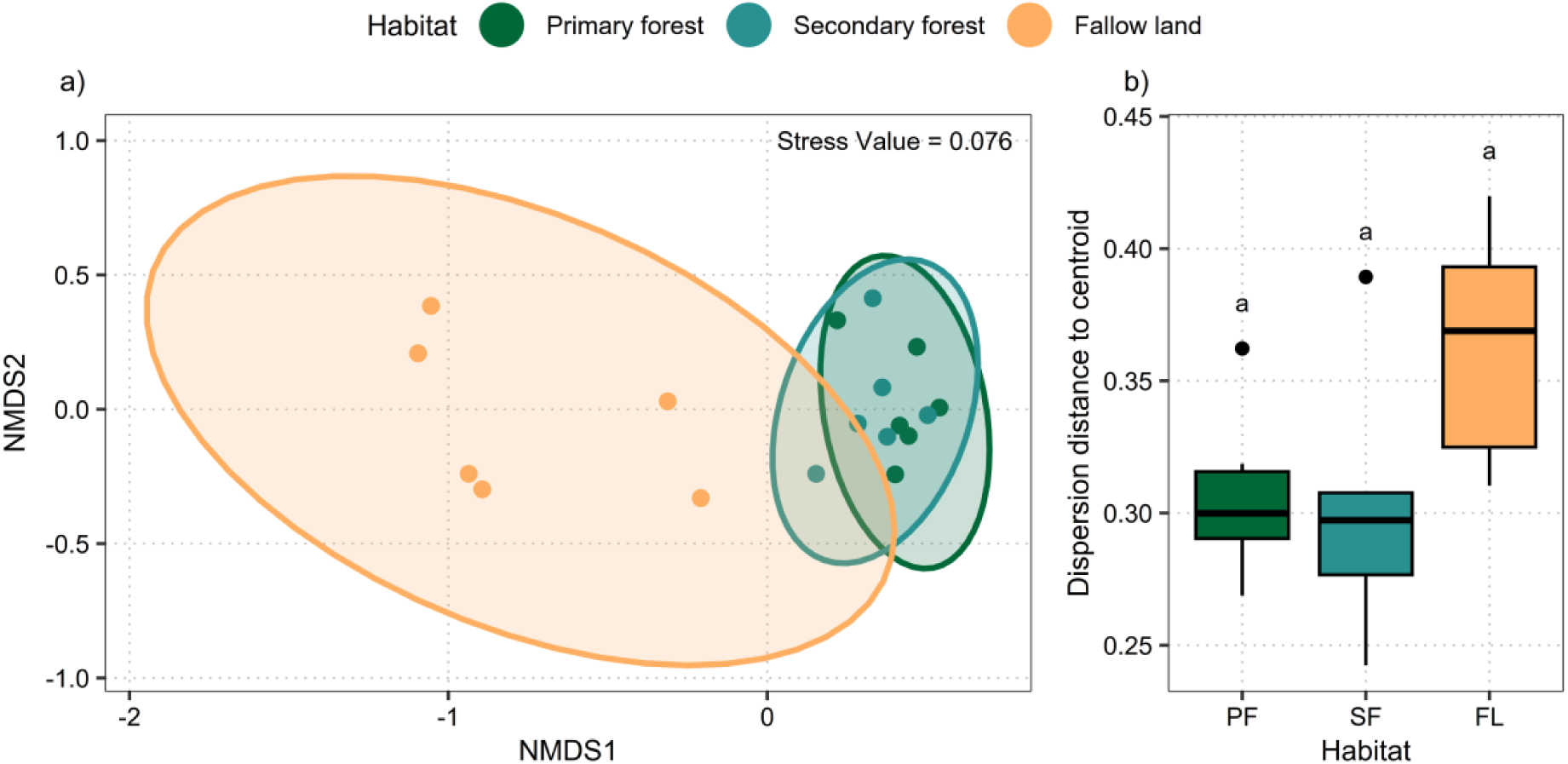
Pattern of species composition in primary forest, secondary forest, and fallow communities. (a) Non-metric Multidimensional Scaling (NMDS) ordination graph representing the distribution of communities from different habitats based on Bray-Curtis dissimilarity distances. (b) Dispersion distance of the different sampled sites to the centroid. The next abbreviations refers to the different habitat types: PF = Primary forest, SF = Secondary forest, and FL = Fallow land. Horizontal lines represent median values, boxes represent the first and third quartiles, and whiskers indicate the maximum and minimum values. Dots represent outliers. Lowercase letters above the boxplots indicate pairwise significant differences between habitats based on Dunn’s test with Holm’s p-value correction.

The overall community rarefaction curve showed a predicted maximum species richness of 197 species (95 % CI: 182 - 222) with a mean log rate constant (lrc) of −1.852 (95 % CI: −2.396 - 1.348) (Figure 5). The sampling performed in the 18 study plots reached a total species richness of 190 species and an estimated species richness of 186 species (95 % CI: 179 - 192; ∼ 94.4 % of the total estimated species richness). Our prediction indicates that a sampling effort of 29 sites would be needed to reach 99 % of species richness detection (95 % CI: 18 - 47).

**Figure 5:**
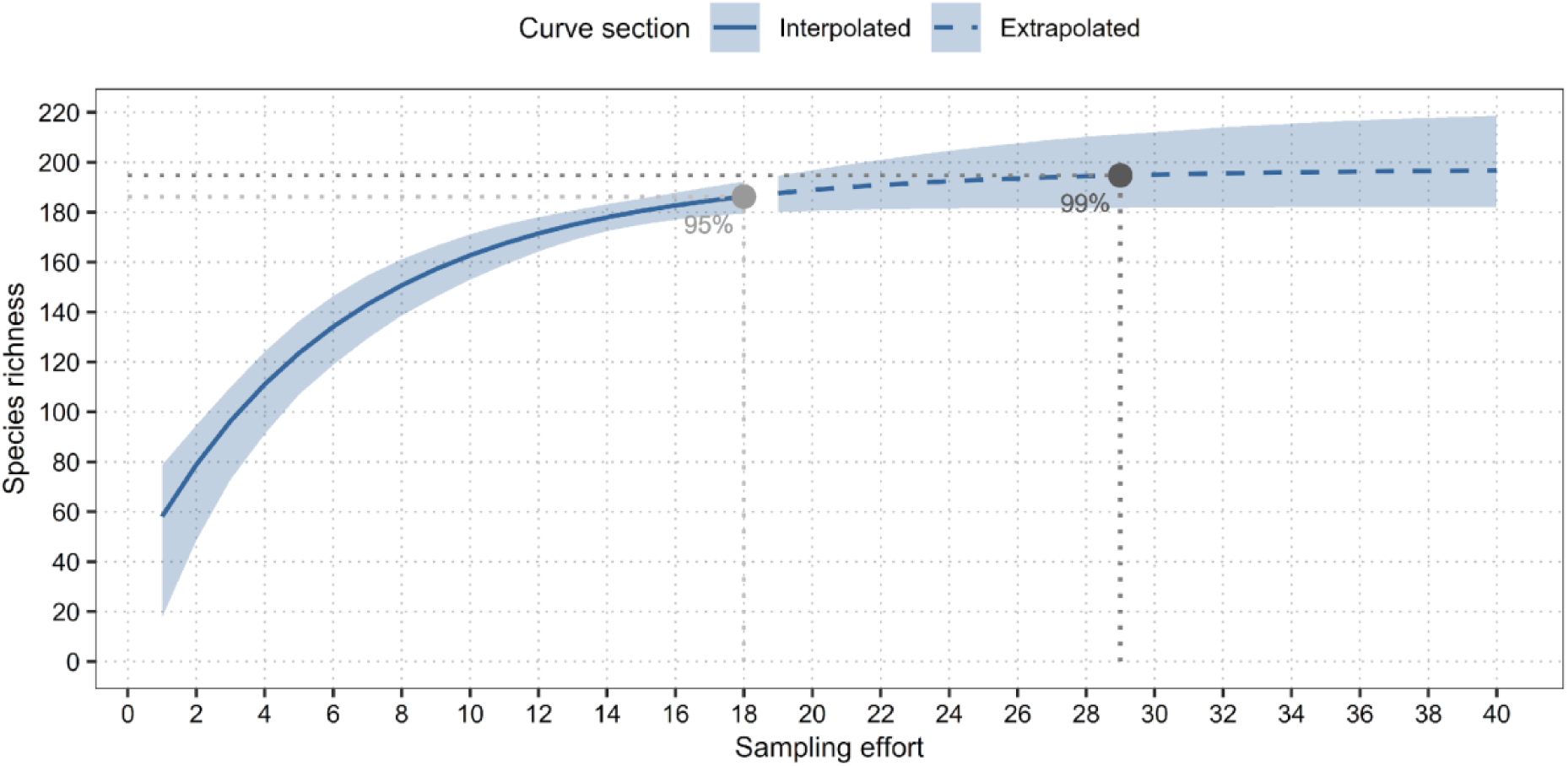
Rarefaction curve of the overall community. The solid line represents the observed species richness, and the dashed line represents the predicted species richness. The shaded area represents the 95% confidence interval of the predicted species richness. A sampling effort unit refers to a sampled plot (four transects of five pitfall traps + five Winkler extractors). Dotted lines represent the sampling effort needed to reach 95% and 99% of the maximum expected species richness.

The rarefaction curves were also estimated separately for each of the different forest types (Figure 6a). The primary forest exhibited the highest predicted maximum species richness with 152 species (95% CI: 144 - 164), followed by the secondary forest with 139 species (95% CI: 131 - 154), and the fallow with 99 species (95% CI: 80 - 198). We also observed differences in the pattern of species accumulation with a steeper slope in the primary forest with a normalized log ratio constant of −0.014 (95 CI: −0.015 - −0.013), followed by the secondary forest with a normalized log ratio constant of −0.015 (95 CI: −0.017 - −0.014) and fallow with a significantly slowest accumulation rate −0.024 (95 CI: −0.028 - 0.019) (Figure 6b).

**Figure 6:**
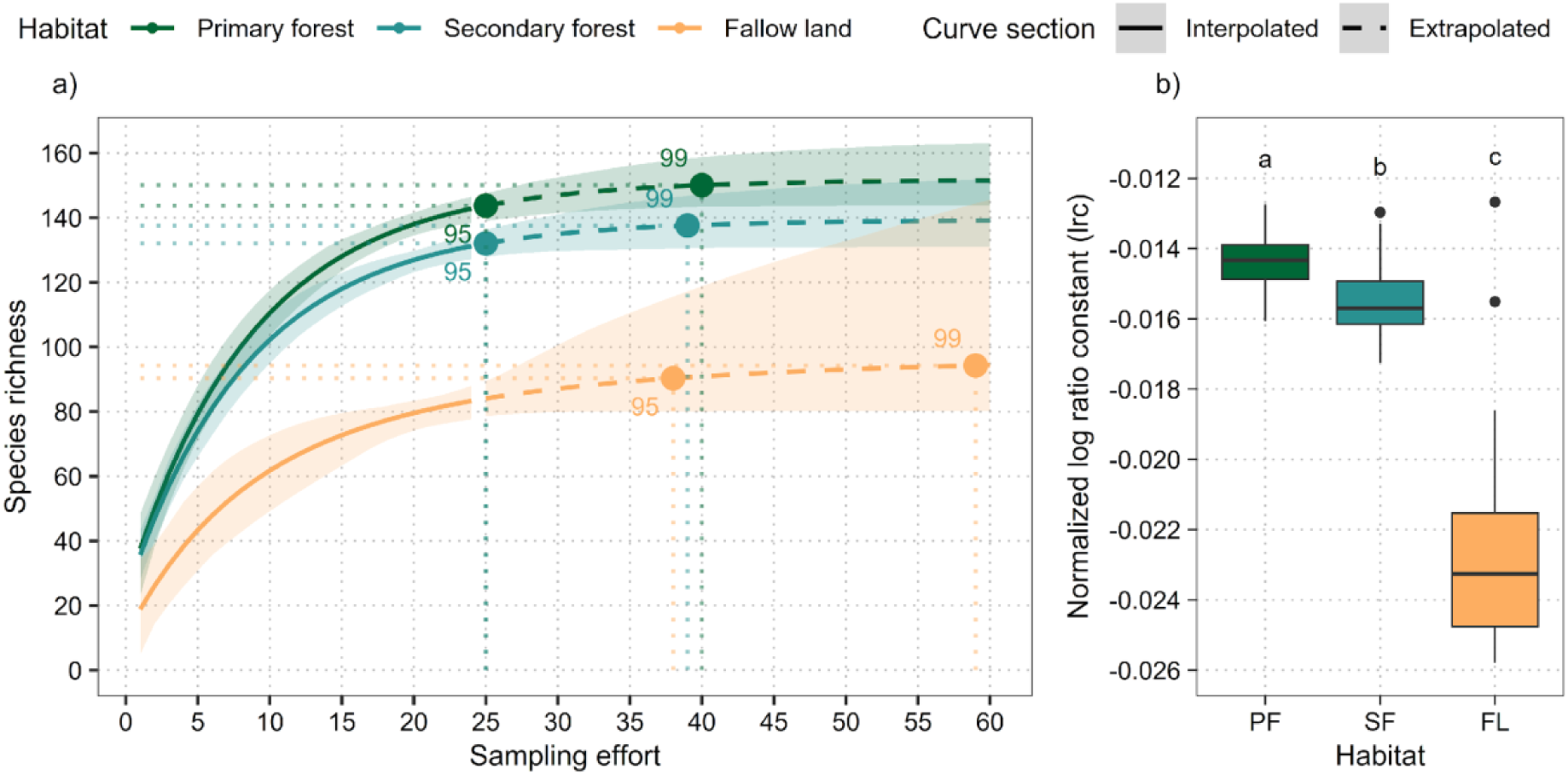
Rarefaction curves and normalized log ratio constant (lrc) values per habitat. (a) Rarefaction curves per habitat. The solid line represents the observed species richness, and the dashed line represents the predicted species richness. The shaded area represents the 95% confidence interval of the predicted species richness. Dotted lines represent the sampling effort (sampling transects) needed to reach 95% and 99% of the maximum expected species richness. (b) Normalized log ratio constant (lrc) values per habitat. The next abbreviations refers to the different habitat types: PF = Primary forest, SF = Secondary forest, and FL = Fallow land. Horizontal lines represent median values, boxes represent the first and third quartiles, and whiskers indicate the maximum and minimum values. Dots represent outliers. Lowercase letters above the boxplots indicate pairwise significant differences between habitats based on Dunn’s test with Holm’s p-value correction (Table S4).

### Pitfall traps versus Winkler extractors

Diversity indices in ant communities differed among sampling method (Kruskal-Wallis test | Species richness (S): d.f. = 5, *χ*^2^ = 24.85, p < 0.001; Shannon index (H’): d.f. = 5, *χ*^2^ = 24.76, p < 0.001; Simpson index: d.f. = 5, *χ*^2^ = 24.73, p < 0.001; Figure 7). Primary and secondary forests presented higher values when winkler extractors were used, but no differences were found in fallow land habitats (Table S5). Moreover, while primary and secondary forests showed higher values compared to fallow land communities for samplings performed with winkler extractors, we could not find differences when we compared the communities sampled with pitfall traps.

**Figure 7:**
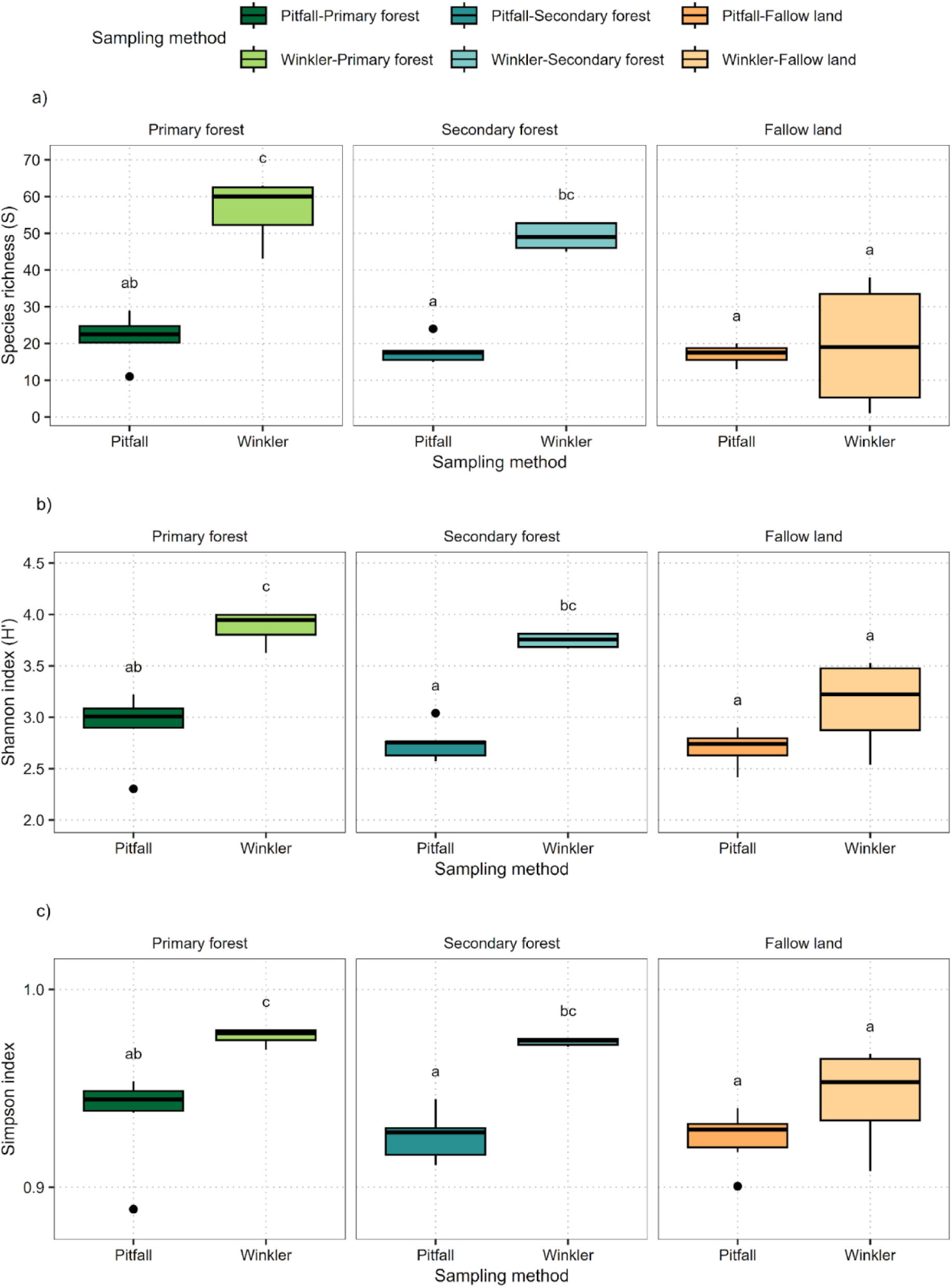
Diversity indices in communities from different habitats sampled with pitfall traps and Winkler extractors. (a) Species richness. (b) Shannon index. and (c) Simpson index. Horizontal lines represent median values, boxes represent the first and third quartiles, and whiskers indicate the maximum and minimum values. Dots represent outliers (extreme outliers were removed to improve visualization). Lowercase letters above the boxplots indicate significant differences between sampling methods and habitat type based on Dunn’s test with Benjamini-Hochberg’s p-value correction. *\*Letters indicate statistical differences between groups*.

**Figure 8.**
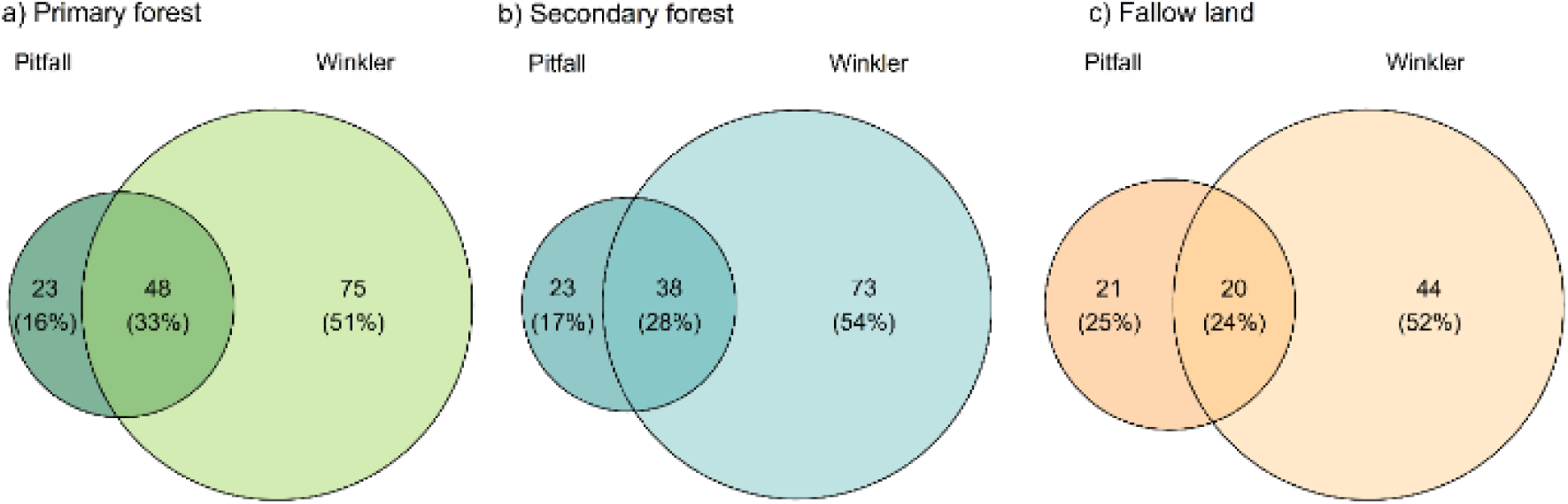
Venn diagram representing the number and percentage of species per habitat and sampling method combination. (a) Primary forest, (b) Secondary forest, and (c) Fallow land. The intersection of the circles represents the number of species shared between the sampling methods, and the non-overlapping areas represent the number of species unique to each sampling method.

In all three habitat types, Winkler extractors captured a greater portion of the litter ant diversity compared to pitfall traps. However, each method also recovered unique species not detected by the other, highlighting their complementarity. This pattern was most pronounced in primary and secondary forests, where Winklers collected the majority of species. In fallow land, although overall diversity was lower, both methods still captured distinct subsets of the community, with limited overlap between them.

Ant communities differed significantly among the sampling method used in the different habitat types (PERMANOVA| habitat: d.f. = 2, F = 5.73, p < 0.001; method: d.f. = 1, F = 12.57, p < 0.001; Figure 9a, Table S6). However, site variability showed a different pattern depending on the habitat type (Figure 9b; Table S7). While the more mature habitat types such as primary and secondary forests showing a higher variability in the communities registered by pitfall traps. The more disturbed fallow land presented a higher dispersion in the communities registered by Winkler extractors.

**Figure 9:**
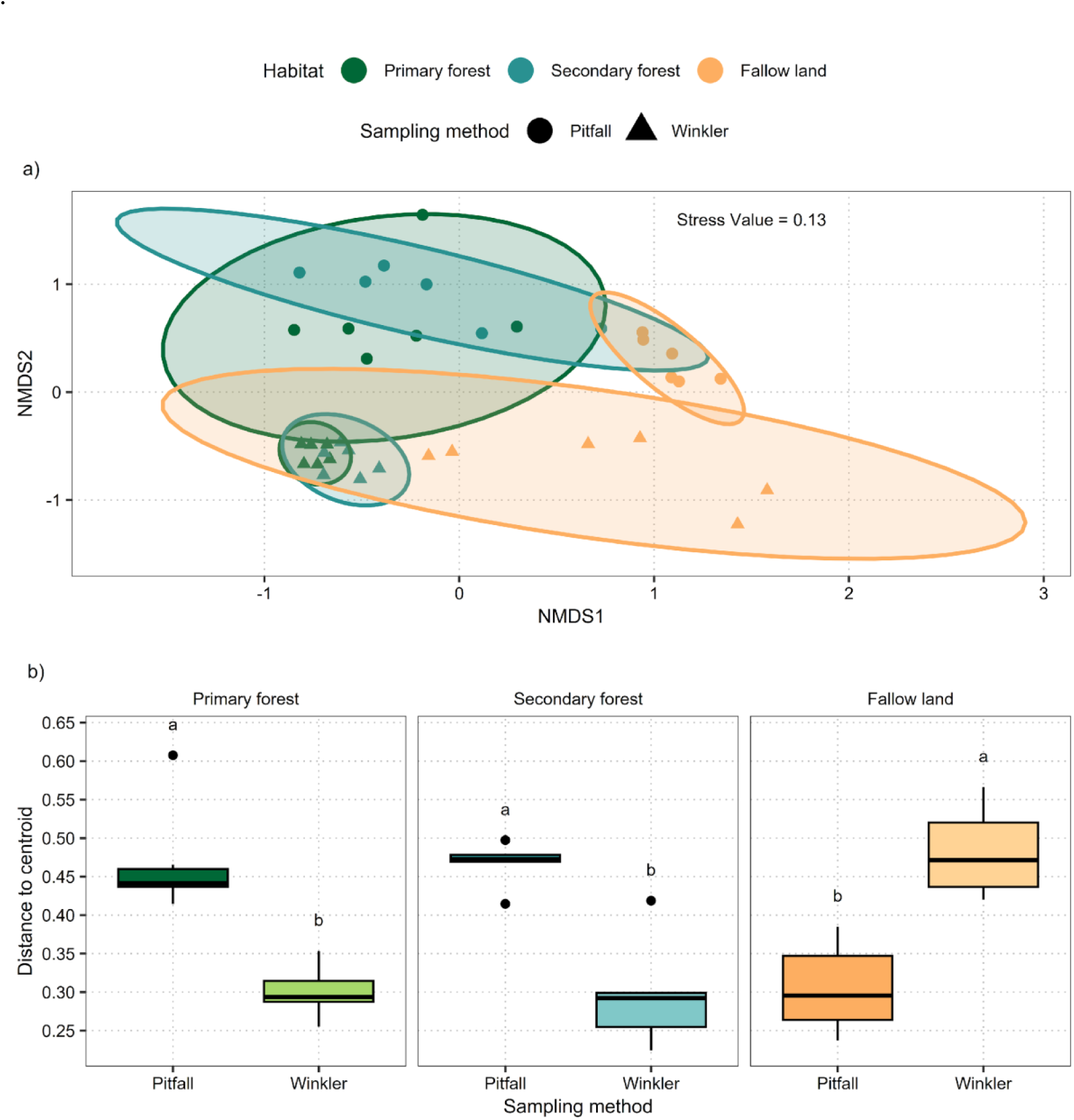
Pattern of species composition in primary forest, secondary forest, and fallow communities according to the sampling method. (a) Non-metric Multidimensional Scaling (NMDS) ordination graph representing the distribution of communities captured with different sampling methods in different habitats based on Bray-Curtis dissimilarity distances. (b) Dispersion distance of the different groups to the centroid. Horizontal lines represent median values, boxes represent the first and third quartiles, and whiskers indicate the maximum and minimum values. Dots represent outliers. Lowercase letters above the boxplots indicate pairwise significant differences between groups based on Dunn’s test with Benjamini-Hochberg’s p-value correction.

The rarefaction curves were also estimated separately for each sampling method (Figure 10a). Pitfall traps showed a predicted maximum species richness of 111 species (95 % CI: 99 - 134). Winkler extractors showed a predicted maximum species richness of 168 species (95 % CI: 151 - 206). We also observed differences in the pattern of species accumulation (Kruskal-Wallis test: d.f. = 1, *χ*^2^ = 1478.95, p < 0.001; Figure 10b) with a more steep slope in the Winkler extractors with a normalized log ratio constant of −0.011 (95 CI: - 0.013 - −0.009), followed by the pitfall traps with a normalized log ratio constant of −0.018 (95 CI: −0.02 - −0.015).

**Figure 10:**
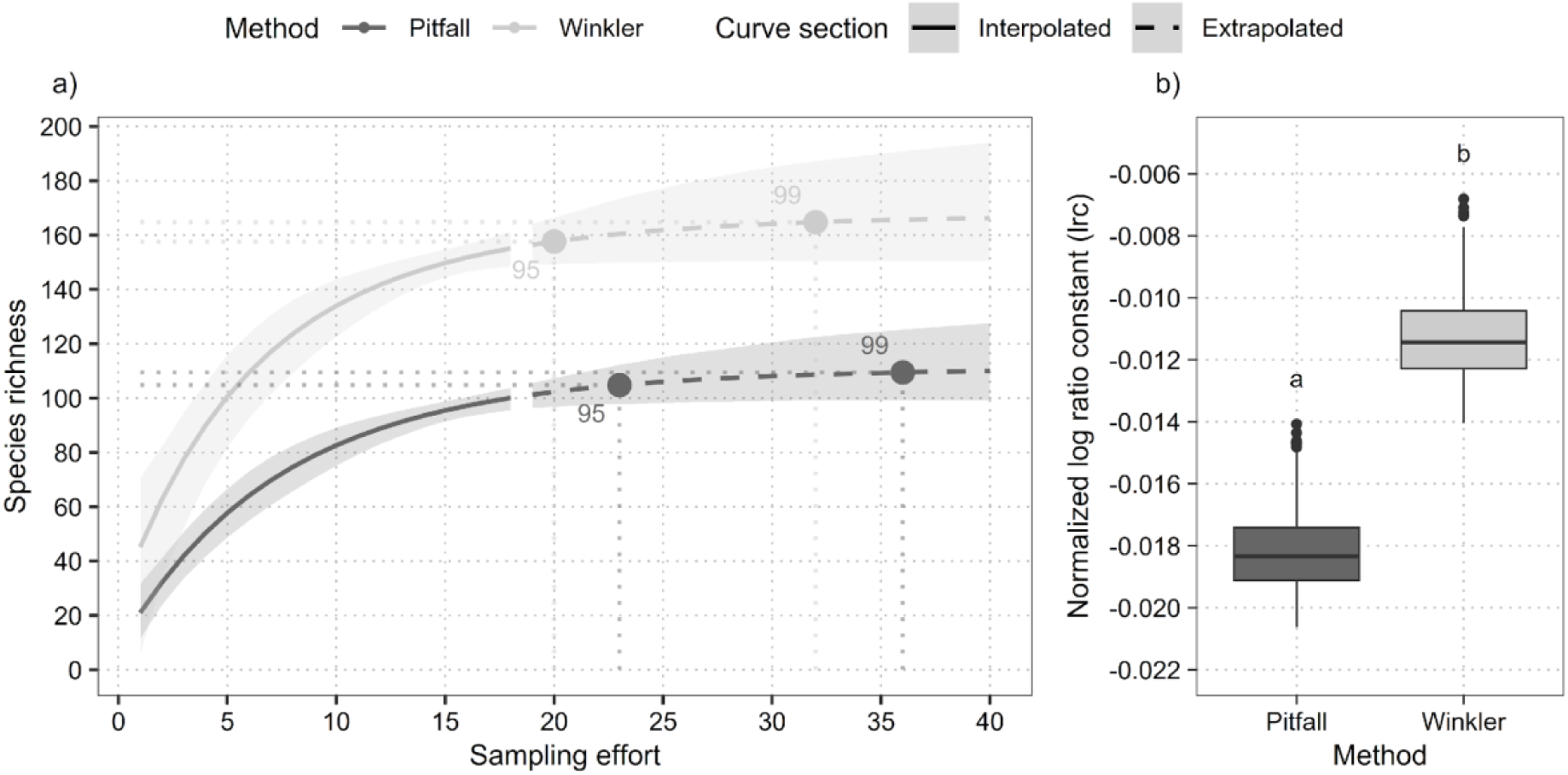
Rarefaction curves and normalized log ratio constant (lrc) values per sampling method. (a) Rarefaction curves per sampling method. The solid line represents the observed species richness, and the dashed line represents the predicted species richness. The shaded area represents the 95% confidence interval of the predicted species richness. Dotted lines represent the sampling effort (sampling plots) needed to reach 95% and 99% of the maximum expected species richness. (b) Normalized log ratio constant (lrc) values per sampling method. Horizontal lines represent median values, boxes represent the first and third quartiles, and whiskers indicate the maximum and minimum values. Dots represent outliers. Lowercase letters above the boxplots indicate pairwise significant differences between habitats based on Kruskal-Wallis’s test.

The sampling performed with pitfall traps reached a total species richness of 102 species and an estimated species richness of 100 species (95 % CI: 96 - 104; ∼ 90.1 % of the total estimated species richness). Our prediction indicates a sampling effort of 36 sampling plots would be needed to reach 99 % of species richness detection (95 % CI: 22 - 64). The sampling performed with Winkler extractors reached a total species richness of 158 species and an estimated species richness of 155 species (95 % CI: 148 - 161; ∼ 92.3 % of the total estimated species richness). Our prediction indicates a sampling effort of 32 sampling plots would be needed to reach 99 % of species richness detection (95 % CI: 17 - 65).

### Pitfall traps versus Winkler extractors in the three different forest habitat types

The rarefaction curves were also separately estimated for each habitat and sampling method combination (Figure 11a, c and d). In primary forest pitfall traps showed a predicted maximum species richness of 85 species (95 % CI: 70 - 118). Winkler extractors showed a predicted maximum species richness of 130 species (95 % CI: 120 - 146). In secondary forest pitfall traps showed a predicted maximum species richness of 78 species (95 % CI: 62 - 127). Winkler extractors showed a predicted maximum species richness of 117 species (95 % CI: 109 - 129). In fallow land pitfall traps showed a predicted maximum species richness of 45 species (95 % CI: 40 - 54). Winkler extractors showed a predicted maximum species richness of 84 species (95 % CI: 62 - 174).

**Figure 11:**
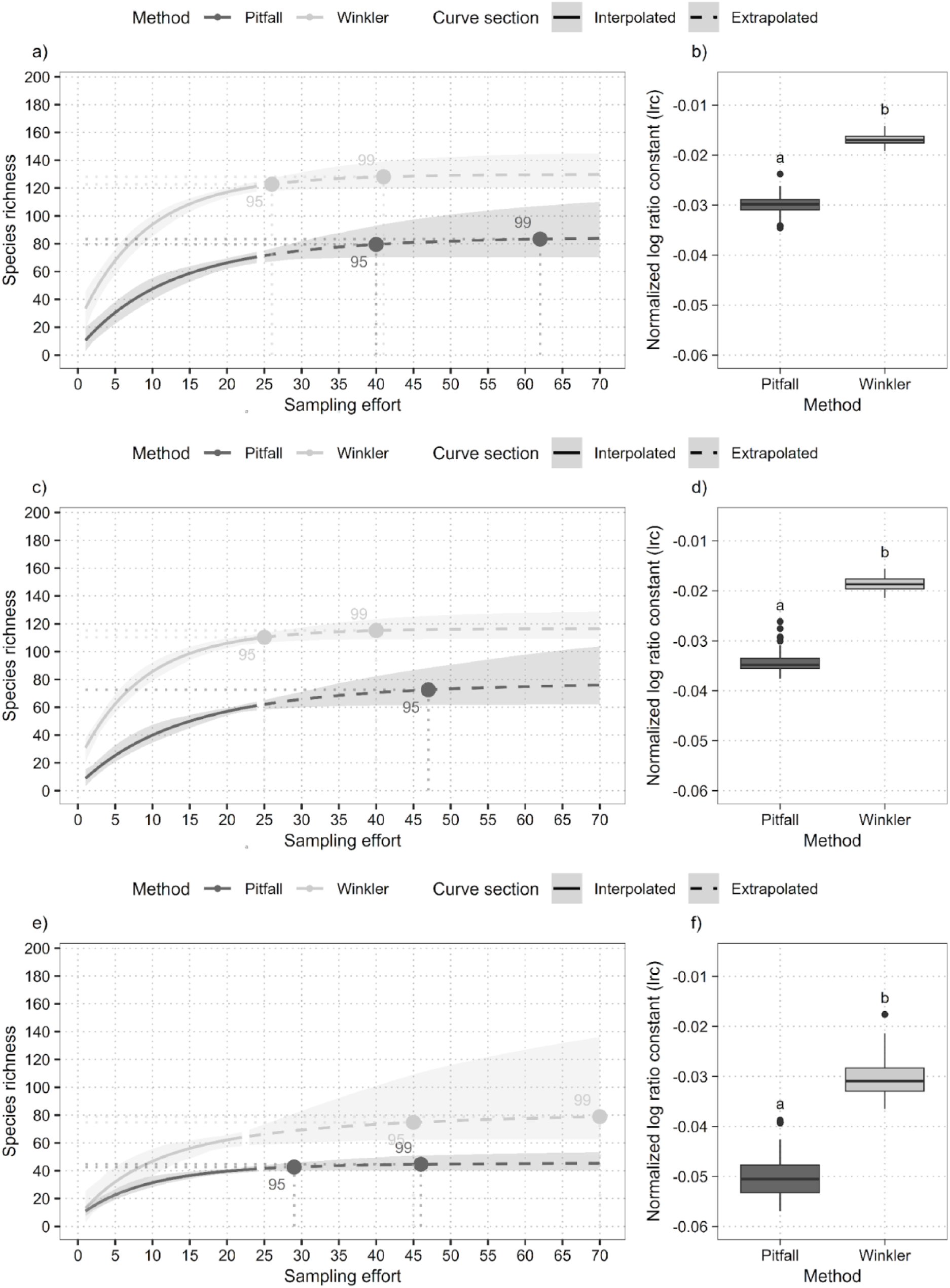
Rarefaction curves and normalized log ratio constant (lrc) values per habitat and sampling method. Rarefaction curves per sampling method for: (a) primary forest, (c) secondary forest, and (e) fallow land. The solid line represents the observed species richness, and the dashed line represents the predicted species richness. The shaded area represents the 95% confidence interval of the predicted species richness. Dotted lines represent the sampling effort (sampling transects) needed to reach 95% and 99% of the maximum expected species richness. Normalized log ratio constant (lrc) values per sampling method for: (b) primary forest, (d) secondary forest, and (f) fallow land. Horizontal lines represent median values, boxes represent the first and third quartiles, and whiskers indicate the maximum and minimum values. Dots represent outliers. Lowercase letters above the boxplots indicate pairwise significant differences between sampling methods per habitat based on Kruskall-Wallis test.

The sampling effort needed to reach 95% of the maximum expected species richness for primary forest with pitfall traps is 40 (95% CI: 20 - 80) and 62 (95% CI: 31 - 124) to reach 99% of the maximum expected species richness. The sampling effort needed to reach 95% of the maximum expected species richness for Winkler extractors in primary forest is 26 (95% CI: 18 - 40), and 41 (95% CI: 28 - 64) to reach 99% of the maximum expected species richness. In secondary forest, the sampling effort needed to reach 95% of the maximum expected species richness for pitfall traps is 47 (95% CI: 21 - 119), and 73 (95% CI: 33 - 183) to reach 99% of the maximum expected species richness. The sampling effort needed to reach 95% of the maximum expected species richness for Winkler extractors in secondary forest is 25 (95% CI: 16 - 38), and 40 (95% CI: 26 - 60) to reach 99% of the maximum expected species richness. In fallow land, the sampling effort needed to reach 95% of the maximum expected species richness for pitfall traps is 29 (95% CI: 14 - 50) and 46 (95% CI: 22 - 79) to reach 99% of the maximum expected species richness. The sampling effort needed to reach 95% of the maximum expected species richness for Winkler extractors in fallow land is 45 (95% CI: 17 - 150), and 70 (95% CI: 26 - 235) to reach 99% of the maximum expected species richness.

The normalized log ratio constant (lrc) value was always higher for Winkler sampling than for pitfall traps in all habitats, indicating that Winkler sampling is a faster method in detecting new species than pitfall traps with a similar sampling effort (Kruskal-Wallis | Primary forest: d.f. = 1, *χ*^2^ = 147, p < 0.001; Secondary forest: d.f. = 1, *χ*^2^ = 147.75, p < 0.001; Fallow land: d.f. = 1, *χ*^2^ = 72, p < 0.001; Fig. 11b, d and f). However, when using Winkler extractors, a greater sampling effort is required in fallow land to reach the maximum expected species richness compared to primary and secondary forests. In contrast, with pitfall traps, higher sampling effort is needed in primary and secondary forests to achieve the same.

## DISCUSSION

### Diversity and ant communities in the different forest types

Our findings show that the diversity and composition of leaf litter ant communities in the Yangambi landscape differ according to forest habitat type. Primary forests exhibited exceptionally high species richness, with a total of 146 ant species recorded, followed by secondary forests with 133 species and 85 species in fallow land. There was no significant difference in species richness between primary and secondary forests, however there was a significant difference in species richness between fallow land and the other two.

The lower species richness of litter ants in fallow land is likely attributable to the habitat degradation, including absence of litter on the soil surface, and limited occurrence of woody plant formations. Moreover, the high species dispersion among sites and the low species richness and accumulation rate observed in fallow land ant communities, suggest that these are a heterogeneous subset of the surrounding mature probably disturbed communities. Their assembly is likely driven by stochastic processes. The low proportion of ant species found exclusively in fallow land may be due to poor plant cover, a lack of litter cover, exposure of the soil surface to sunlight and the absence of microhabitats (e.g. scarcity of dead wood, scarcity of large trees and scarcity of termite mounds) (Devineau 1975; Kaspari et al. 2000). These factors sometimes limit the colonisation and survival of many species, particularly those that are habitat specialists or dependent on a complex forest environment (Delabie and Fowler 1995; Bastos and Harada 2011). It is likely that this pattern extends to other degraded sites within the DRC. Moreover, primary forests, despite being more diverse in plant species composition, harbour more homogeneous communities. The heterogeneity detected in fallow lands suggests that these disturbed habitats may require more extensive sampling than other habitats to capture the full diversity of ant communities (see also Fimbel et al. 2001; Daily et al. 2001; Kouakou 2009).

Our results provide clear evidence that ecological disturbance in the Yangambi landscape has strongly contributed to a reduction in the abundance and species richness of fallow litter ants and underline the importance of preserving primary forests and sustainably managing secondary forests to limit the erosion of biodiversity and promote ecosystem resilience. In addition, most of the ants in the two forest habitats are cryptic species that live in the litter and rotting trunks (mainly *Strumigenys*, *Parasycia* and *Anochetus*), whereas the species in the fallow mainly nest underground because annual fires burn the litter and other dead wood above ground. Underground ant assemblages tend to be less species-rich than aboveground assemblages (Yeo et al. 2017).

Other studies have shown that heavy rainfall during the wet season associated with sloping rocky ground can prevent the creation of nests by certain ant species in areas where vegetation is scarce and that these species very often create their nests in areas where this disturbance is minimal (Majiner and Delabie 1994; Mertl et al. 2009). Hence fallow land, although lacking significant woody cover, facilitates the habitation of certain ant species that are very often generalists and like open spaces, for example the majority of Formicinae and Dolichoderinae (Kouakou 2009).

### Sampling methods

As in other studies on leaf litter ants in tropical regions, Winkler extractors yielded higher species diversity than pitfall traps (Fisher et al. 2000; Lopes and Vasconcelos 2008; Orsolon-Souza et al. 2011), though the latter contributed complementary species to the overall community. Pitfall traps appear to capture species active during different time periods—such as nocturnal fauna—that may be absent from the litter at the time it is collected for Winkler extraction (Martelli et al. 2004; Orsolon-Souza et al. 2011; Sabu et al. 2011). Kaspari (2000), suggests that the Winkler technique can only be applied in vegetation covering a thick layer of litter making it possible to capture entomofauna that are not very mobile and that depend on the soil’s litter layer. It is often concluded that these two methods should always be used together during leaf litter ant surveys (York 2000; Martelli et al. 2004; Silva et al. 2013). Winkler sampling is a faster method in detecting new species than pitfall traps with a similar sampling effort. The effectiveness of the Winkler method in this study is evident from its frequent success in capturing the small-bodied leaf litter ants (Agosti and Alonso 2000; York 2000; Agosti and Alonso 2003). As an example of this, we recorded 25 species of the cryptic genus *Strumigenys* during this study and all of these species were captured using the Winkler method, with only 4 species also captured at the same time using the Pitfall method. The majority of the *Strumigenys* species are small and nest and feed in leaf litter, soil or rotting wood, while only a few species prefer an arboreal lifestyle (Bolton 2000). Moreover, all the Amblyoponinae and Proceratiinae species were captured exclusively using the Winkler method. Very often Amblyoponinae and Proceratiinae species are rarely observed on the ground surface as they nest in the soil, under wood, in logs or in rotten wood on the ground (Yoshimura et al. 2012; Blaimer et al. 2015; Eguchi et al. 2015; Ward and Fisher 2016).

### Main Conclusions and Implications for Future Research

This study demonstrates that vegetation disturbance reduces ant diversity, while forest recovery enhances invertebrate richness. Future research in the Yangambi reserve should include sampling the arboreal ant fauna, which typically harbours one third of the tropical fauna (Philpott and Armbrecht, 2006, Leponce et al. 2019, Yodé et al. 2020). Also, an interesting addition would be sampling primary rainforest plots further remote in the forest, to analyse the influence of fragmentation. In addition, the same study should be repeated at other sites in order to gain a broader understanding of the diversity of ants in the different vegetation in the tropical region of the Congo Basin and particularly in the DRC.

## ACKNOWLEDGEMENTS

We want to thank the staff of CEBioS, Capacities for Biodiversity and Sustainable Development, a programme financed by the Directorate-General for Development Cooperation (DGD) and housed at the Royal Belgian Institute of Natural Sciences (RBINS), where it belongs to the Operational Directorate ‘Natural Environment’ and more specifically the BioPolS group (Belgian Biodiversity Policy Support Group) for their continuously support for Afrotropical myrmecological studies during the last 15 years. Moreover, we are in debt to one anonymous referee for suggestions on an earlier version of the manuscript. We also want to thank Benjamin Toirambe Bamoninga, Autorité Nationale Compétente APA/RDC du Ministère de l’Environnement et Développement Durable: Permis N008/SG-EDD/BTB/ANCCB-RDC/04/2024.

## Appendix

### Sampling detail

*F=Fallow ; PS=Primary forest ; Secondary forest ; GT= Grand total ; TP=Type of traps ; W=Winkler ; P= Pitfalls*

**Table 1.** Sampling detail.

| Subfamily and species | F | PS | SF | GT | TP |
| --- | --- | --- | --- | --- | --- |
| **Amblyoponinae** | **0** | **9** | **7** | **16** |  |
| Prionopelta amieti Terron, 1974 | 0 | 7 | 5 | 12 | W |
| Stigmatomma kgac-drc01 | 0 | 2 | 0 | 2 | W |
| Stigmatomma kgac-drc02 | 0 | 0 | 2 | 2 | W |
| **Dolichoderinae** | **26** | **51** | **48** | **125** |  |
| Tapinoma kgac-drc01 | 5 | 1 | 0 | 6 | PW |
| Tapinoma kgac-drc02 | 9 | 1 | 0 | 10 | PW |
| Technomyrmex andrei Emery, 1899 | 1 | 28 | 19 | 48 | PW |
| Technomyrmex laurenti (Emery, 1899) | 0 | 6 | 7 | 13 | PW |
| Technomyrmex lujae (Forel, 1905) | 4 | 7 | 9 | 20 | PW |
| Technomyrmex moerens Santschi, 1913 | 4 | 3 | 1 | 8 | PW |
| Technomyrmex nigriventris Santschi, 1910 | 0 | 0 | 2 | 2 | W |
| Technomyrmex parviflavus Bolton, 2007 | 3 | 5 | 10 | 18 | W |
| **Dorylinae** | **4** | **26** | **25** | **55** |  |
| Aenictus bidentatus Donisthorpe, 1942 | 0 | 1 | 0 | 1 | W |
| Dorylus kgac-drc01 | 1 | 6 | 4 | 11 | PW |
| Dorylus kgac-drc02 | 0 | 0 | 1 | 1 | P |
| Dorylus kgac-drc03 | 0 | 5 | 7 | 12 | PW |
| Dorylus kgac-drc04 | 0 | 2 | 1 | 3 | P |
| Dorylus kgac-drc05 | 0 | 0 | 1 | 1 | P |
| Dorylus kgac-drc06 | 0 | 1 | 0 | 1 | P |
| Dorylus wilverthi Emery, 1899 | 3 | 0 | 0 | 3 | P |
| Eburopone wroughtoni (Forel, 1910) | 0 | 2 | 0 | 2 | W |
| Lioponera foreli (Santschi, 1914) | 0 | 1 | 1 | 2 | W |
| Parasyscia afr-par15 | 0 | 1 | 0 | 1 | W |
| Parasyscia kgac-drc01 | 0 | 2 | 0 | 2 | W |
| Parasyscia kgac-drc02 | 0 | 2 | 0 | 2 | PW |
| Parasyscia nitidulus (Brown, 1975) | 0 | 3 | 10 | 13 | W |
| **Formicinae** | **139** | **52** | **62** | **253** |  |
| Anoplolepis kohli (Forel, 1916) | 0 | 7 | 10 | 17 | PW |
| Camponotus acvapimensis Mayr, 1862 | 24 | 1 | 3 | 28 | PW |
| Camponotus caesar imperator Emery, 1899 | 3 | 2 | 9 | 14 | P |
| Camponotus chrysurus Gerstäcker, 1871 | 0 | 1 | 1 | 2 | P |
| Camponotus crawleyi Emery, 1920 | 13 | 16 | 19 | 48 | PW |
| Camponotus foraminosus Forel, 1879 | 12 | 1 | 0 | 13 | P |
| Camponotus guttatus Emery, 1899 | 0 | 1 | 2 | 3 | P |
| Camponotus kgac-drc01 | 0 | 1 | 0 | 1 | W |
| Camponotus kgac-drc02 | 18 | 0 | 2 | 20 | P |
| Camponotus schoutedeni Forel, 1911 | 5 | 0 | 0 | 5 | P |
| Lepisiota capensis anceps (Forel, 1916) | 20 | 1 | 0 | 21 | PW |
| Lepisiota kgac-drc01 | 10 | 0 | 0 | 10 | WP |
| Lepisiota kgac-drc02 | 1 | 0 | 0 | 1 | P |
| Nylanderia jaegerskioeldi (Mayr, 1904) | 25 | 2 | 2 | 29 | PW |
| Nylanderia lepida (Santschi, 1915) | 0 | 12 | 4 | 16 | PW |
| Nylanderia umbella LaPolla & Fisher, 2011 | 0 | 1 | 0 | 1 | W |
| Oecophylla longinoda (Latreille, 1802) | 1 | 1 | 1 | 3 | PW |
| Oecophylla longinoda fusca Emery, 1899 | 0 | 0 | 1 | 1 | P |
| Paraparatrechina concinnata LaPolla & Cheng, 2010 | 0 | 0 | 1 | 1 | P |
| Paraparatrechina weissi (Santschi, 1910) | 2 | 3 | 1 | 6 | W |
| Paratrechina longicornis (Latreille, 1802) | 1 | 2 | 1 | 4 | PW |
| Plagiolepis kgac-drc01 | 1 | 0 | 0 | 1 | W |
| Polyrhachis militaris (Fabricius, 1782) | 3 | 0 | 5 | 8 | P |
| **Myrmicinae** | **210** | **563** | **500** | **1273** |  |
| Baracidris sitra Bolton, 1981 | 0 | 4 | 0 | 4 | W |
| Calyptomyrmex brevis Weber, 1943 | 0 | 4 | 5 | 9 | W |
| Cardiocondyla emeryi Forel, 1881 | 5 | 0 | 0 | 5 | PW |
| Carebara distincta (Bolton & Belshaw, 1993) | 3 | 4 | 3 | 10 | W |
| Carebara kgac-drc01 | 6 | 24 | 21 | 51 | PW |
| Carebara kgac-drc02 | 9 | 9 | 11 | 29 | PW |
| Carebara kgac-drc03 | 0 | 9 | 12 | 21 | PW |
| Carebara kgac-drc04 | 1 | 0 | 0 | 1 | W |
| Carebara kgac-drc05 | 0 | 5 | 4 | 9 | W |
| Carebara kgac-drc06 | 0 | 1 | 2 | 3 | W |
| Carebara kgac-drc07 | 0 | 0 | 1 | 1 | W |
| Carebara kgac-drc08 | 0 | 0 | 1 | 1 | W |
| Carebara kgac-drc09 | 0 | 0 | 1 | 1 | W |
| Carebara silvestrii (Santschi, 1914) | 2 | 7 | 5 | 14 | PW |
| Carebara villiersi (Bernard, 1953) | 0 | 0 | 2 | 2 | W |
| Cataulacus erinaceus Stitz, 1910 | 1 | 1 | 1 | 3 | P |
| Cataulacus guineensis Smith, 1853 | 2 | 1 | 3 | 6 | PW |
| Cataulacus traegaardhi Santschi, 1914 | 3 | 0 | 0 | 3 | PW |
| Crematogaster africana Mayr, 1895 | 0 | 2 | 5 | 7 | PW |
| Crematogaster clariventris Mayr, 1895 | 0 | 0 | 1 | 1 | P |
| Crematogaster kgac-drc01 | 0 | 5 | 7 | 12 | PW |
| Crematogaster kgac-drc02 | 0 | 2 | 1 | 3 | P |
| Crematogaster kgac-drc03 | 2 | 14 | 13 | 29 | PW |
| Crematogaster kgac-drc04 | 1 | 1 | 3 | 5 | PW |
| Crematogaster kgac-drc05 | 0 | 0 | 1 | 1 | P |
| Crematogaster kgac-drc06 | 0 | 1 | 0 | 1 | W |
| Crematogaster kgac-drc07 | 1 | 0 | 0 | 1 | P |
| Crematogaster kgac-drc08 | 0 | 2 | 0 | 2 | P |
| Crematogaster kgac-drc09 | 0 | 0 | 1 | 1 | P |
| Crematogaster kgac-drc10 | 0 | 0 | 1 | 1 | W |
| Crematogaster stadelmanni Mayr, 1895 | 0 | 1 | 1 | 2 | PW |
| Crematogaster wellmani Forel, 1909 | 0 | 2 | 0 | 2 | PW |
| Dicroaspis laevidens (Santschi, 1919) | 0 | 1 | 1 | 2 | W |
| Meranoplus inermis Emery, 1895 | 1 | 0 | 0 | 1 | W |
| Microdacetone tibialis Weber, 1952 | 0 | 0 | 1 | 1 | W |
| Monomorium exiguum Forel, 1894 | 13 | 0 | 1 | 14 | PW |
| Monomorium invidium Bolton, 1987 | 13 | 21 | 21 | 55 | PW |
| Monomorium kgac-drc02 | 0 | 0 | 1 | 1 | W |
| Monomorium kgac-drc03 | 1 | 0 | 0 | 1 | W |
| Monomorium madecassum Forel, 1892 | 0 | 0 | 2 | 2 | W |
| Monomorium pharaonis (Linnaeus, 1758) | 7 | 9 | 9 | 25 | PW |
| Monomorium rosae Santschi, 1920 | 1 | 0 | 0 | 1 | W |
| Monomorium rotundatum Santschi, 1920 | 1 | 1 | 4 | 6 | W |
| Monomorium strangulatum Santschi, 1921 | 1 | 3 | 5 | 9 | PW |
| Monomorium termitobium Forel, 1892 | 0 | 0 | 3 | 3 | W |
| Myrmecaria sp01 | 2 | 15 | 11 | 28 | PW |
| Nesomyrmex evelynae (Forel, 1916) | 0 | 1 | 0 | 1 | W |
| Pheidole batrachorum Wheeler, 1922 | 0 | 4 | 2 | 6 | PW |
| Pheidole buchholzi Mayr, 1901 | 0 | 19 | 4 | 23 | PW |
| Pheidole glabrella Fischer, Hita Garcia & Peters, 2012 | 0 | 5 | 2 | 7 | PW |
| Pheidole kgac-drc01 | 0 | 8 | 5 | 13 | PW |
| Pheidole kgac-drc02 | 4 | 4 | 2 | 10 | PW |
| Pheidole kgac-drc03 | 2 | 13 | 6 | 21 | PW |
| Pheidole kgac-drc04 | 11 | 23 | 20 | 54 | PW |
| Pheidole kgac-drc05 | 1 | 14 | 12 | 27 | PW |
| Pheidole kgac-drc06 | 4 | 7 | 6 | 17 | PW |
| Pheidole megacephala (Fabricius, 1793) | 36 | 5 | 13 | 54 | PW |
| Pheidole pulchella Santschi, 1910 | 0 | 1 | 0 | 1 | P |
| Pristomyrmex africanus Karavaiev, 1931 | 1 | 17 | 5 | 23 | W |
| Pristomyrmex orbiceps (Santschi, 1914) | 3 | 20 | 20 | 43 | PW |
| Strumigenys bellatrix (Bolton, 2000) | 0 | 4 | 6 | 10 | W |
| Strumigenys bernardi Brown, 1960 | 6 | 22 | 24 | 52 | PW |
| Strumigenys bitheria Bolton, 1983 | 2 | 16 | 13 | 31 | W |
| Strumigenys cavinasis (Brown, 1950) | 1 | 2 | 1 | 4 | W |
| Strumigenys concolor Santschi, 1914 | 4 | 17 | 23 | 44 | PW |
| Strumigenys dotaja (Bolton, 1983) | 1 | 11 | 8 | 20 | PW |
| Strumigenys enkara (Bolton, 1983) | 0 | 1 | 2 | 3 | W |
| Strumigenys hensekta (Bolton, 1983) | 0 | 3 | 4 | 7 | W |
| Strumigenys kersa (Bolton, 1983) | 0 | 1 | 0 | 1 | W |
| Strumigenys kgac-drc01 | 0 | 5 | 5 | 10 | W |
| Strumigenys kgac-drc02 | 0 | 2 | 3 | 5 | W |
| Strumigenys kgac-drc03 | 1 | 6 | 7 | 14 | W |
| Strumigenys ludovici Forel, 1904 | 0 | 1 | 0 | 1 | W |
| Strumigenys lujae Forel, 1902 | 0 | 5 | 1 | 6 | W |
| Strumigenys marginata (Santschi, 1914) | 0 | 1 | 0 | 1 | W |
| Strumigenys ninda (Bolton, 1983) | 0 | 13 | 14 | 27 | W |
| Strumigenys petiolata Bernard, 1953 | 5 | 2 | 6 | 13 | W |
| Strumigenys ravidura (Bolton, 1983) | 0 | 1 | 4 | 5 | W |
| Strumigenys rogeri Emery, 1890 | 6 | 2 | 1 | 9 | W |
| Strumigenys roomi (Bolton, 1972) | 1 | 0 | 2 | 3 | W |
| Strumigenys serrula Santschi, 1910 | 5 | 17 | 18 | 40 | PW |
| Strumigenys sharra (Bolton, 1983) | 0 | 1 | 0 | 1 | W |
| Strumigenys tacta (Bolton, 1983) | 0 | 12 | 6 | 18 | W |
| Strumigenys tetragnatha (Taylor, 1966) | 0 | 1 | 0 | 1 | W |
| Strumigenys tetraphanes Brown, 1954 | 0 | 5 | 4 | 9 | W |
| Sylophopsis cryptobia Santschi, 1921 | 8 | 17 | 21 | 46 | PW |
| Sylophopsis malamixta (Bolton, 1987) | 0 | 1 | 0 | 1 | W |
| Tetramorium ataxium Bolton, 1980 | 1 | 0 | 0 | 1 | P |
| Tetramorium coloreum Mayr, 1901 | 0 | 4 | 1 | 5 | PW |
| Tetramorium jugatum Bolton, 1980 | 0 | 5 | 0 | 5 | PW |
| Tetramorium kgac-drc01 | 0 | 0 | 1 | 1 | W |
| Tetramorium kgac-drc02 | 3 | 12 | 4 | 19 | PW |
| Tetramorium kgac-drc03 | 0 | 0 | 1 | 1 | W |
| Tetramorium kgac-drc04 | 0 | 1 | 0 | 1 | P |
| Tetramorium metactum Bolton, 1980 | 0 | 1 | 0 | 1 | W |
| Tetramorium murali Forel, 1910 | 0 | 1 | 1 | 2 | W |
| Tetramorium muscorum Arnold, 1926 | 4 | 15 | 14 | 33 | PW |
| Tetramorium peutli Forel, 1916 | 0 | 6 | 4 | 10 | W |
| Tetramorium psyanum Bolton, 1980 | 0 | 1 | 0 | 1 | W |
| Tetramorium pullulum Santschi, 1924 | 0 | 1 | 0 | 1 | W |
| Tetramorium sericeiventris Emery, 1877 | 9 | 0 | 1 | 10 | PW |
| Tetramorium simillimum (Smith, 1851) | 13 | 18 | 12 | 43 | PW |
| Tetramorium snellingi Hita Garcia, Fischer & Peters, 2010 | 0 | 2 | 0 | 2 | W |
| Tetramorium susannae Hita Garcia, Fischer & Peters, 2010 | 0 | 14 | 3 | 17 | PW |
| Tetramorium zambeziense Santschi, 1939 | 2 | 23 | 21 | 46 | PW |
| **Ponerinae** | **43** | **147** | **133** | **323** |  |
| Anochetus africanus (Mayr, 1865) | 2 | 9 | 4 | 15 | W |
| Anochetus katonae Forel, 1907 | 0 | 7 | 5 | 12 | W |
| Anochetus kgac-drc01 | 0 | 2 | 1 | 3 | W |
| Anochetus kgac-drc02 | 0 | 1 | 0 | 1 | W |
| Anochetus kgac-drc03 | 0 | 0 | 1 | 1 | P |
| Bothroponera ancilla (Emery, 1899) | 0 | 1 | 0 | 1 | W |
| Bothroponera kgac-drc01 | 0 | 1 | 0 | 1 | W |
| Bothroponera pachyderma (Emery, 1901) | 0 | 5 | 3 | 8 | PW |
| Bothroponera rubescens Santschi, 1937 | 1 | 6 | 5 | 12 | W |
| Bothroponera soror (Emery, 1899) | 2 | 6 | 7 | 15 | PW |
| Bothroponera talpa André, 1890 | 0 | 3 | 1 | 4 | W |
| Euponera bruno (Forel, 1913) | 1 | 1 | 1 | 3 | W |
| Fisheropone ambigua (Weber, 1942) | 0 | 2 | 0 | 2 | W |
| Hypoponera angustata (Santschi, 1914) | 2 | 0 | 3 | 5 | W |
| Hypoponera inaudax (Santschi, 1919) | 0 | 4 | 1 | 5 | W |
| Hypoponera kgac-drc01 | 3 | 22 | 23 | 48 | PW |
| Hypoponera molesta Bolton & Fisher, 2011 | 1 | 9 | 7 | 17 | W |
| Hypoponera perparva Bolton & Fisher, 2011 | 2 | 21 | 22 | 45 | W |
| Leptogenys camerunensis Stitz, 1910 | 0 | 0 | 1 | 1 | P |
| Leptogenys crustosa Santschi, 1914 | 1 | 0 | 0 | 1 | P |
| Megaponera analis (Latreille, 1802) | 3 | 2 | 2 | 7 | PW |
| Mesoponera ambigua (André, 1890) | 6 | 13 | 16 | 35 | W |
| Mesoponera ingesta (Wheeler, 1922) | 4 | 9 | 10 | 23 | PW |
| Mesoponera kgac-drc01 | 0 | 7 | 3 | 10 | WP |
| Mesoponera kgac-drc02 | 0 | 0 | 1 | 1 | P |
| Odontomachus assiniensis Emery, 1892 | 0 | 5 | 2 | 7 | WP |
| Odontomachus troglodytes Santschi, 1914 | 14 | 5 | 13 | 32 | PW |
| Paltothyreus tarsatus (Fabricius, 1798) | 0 | 2 | 0 | 2 | PW |
| Phrynoponera bequaerti Wheeler, 1922 | 1 | 2 | 1 | 4 | W |
| Phrynoponera gabonensis (André, 1892) | 0 | 1 | 0 | 1 | W |
| Platythyrea gracillima Wheeler, 1922 | 0 | 1 | 0 | 1 | PW |
| **Proceratiinae** | **1** | **12** | **4** | **17** |  |
| Discothyrea mixta Brown, 1958 | 0 | 7 | 0 | 7 | W |
| Discothyrea venus Hita-Garcia & Lieberman, 2019 | 1 | 4 | 4 | 9 | W |
| Probolomyrmex guineensis Taylor, 1965 | 0 | 1 | 0 | 1 | W |
| **Pseudomyrmecinae** | **1** | **4** | **0** | **5** |  |
| Tetraponera aethiops Smith, 1877 | 0 | 3 | 0 | 3 | PW |
| Tetraponera anthracina (Santschi, 1910) | 1 | 0 | 0 | 1 | P |
| Tetraponera latifrons (Emery, 1912) | 0 | 1 | 0 | 1 | P |
| **Grand total** | **424** | **864** | **779** | **2067** |  |

## Supporting information

**Table S1:** Pairwise Dunn’s test results for the comparison of the effect of habitat type on diversity indices. (a) Species richness (S), (b) Shannon index (H), and (c) Simpson index.

a) Species richness
| Pair comparisons | Z | p |
| --- | --- | --- |
| Primary forest - Secondary forest | 1.14 | 0.255 |
| Primary forest - Fallow land | -3.49 | 0.001** |
| Secondary forest - Fallow land | -2.36 | 0.037* |

\* $p \leq 0.05$ , \*\* $p \leq 0.01$ , \*\*\* $p \leq 0.001$

b) Shannon index
| Pair comparisons | Z | p |
| --- | --- | --- |
| Primary forest - Secondary forest | 1.19 | 0.234 |
| Primary forest - Fallow land | -3.51 | 0.001** |
| Secondary forest - Fallow land | -2.33 | 0.040* |

\* $p \leq 0.05$ , \*\* $p \leq 0.01$ , \*\*\* $p \leq 0.001$

c) Simpson index
| Pair comparisons | Z | p |
| --- | --- | --- |
| Primary forest - Secondary forest | 1.08 | 0.279 |
| Primary forest - Fallow land | -3.46 | 0.002** |
| Secondary forest - Fallow land | -2.38 | 0.035* |

\* $p \leq 0.05$ , \*\* $p \leq 0.01$ , \*\*\* $p \leq 0.001$

**Table S2:** Permanova pairwise test results for the comparison of the effect of habitat type on the community composition of ants.

| Pair comparisons | d.f. | F | p |
| --- | --- | --- | --- |
| Primary forest vs Fallow land | 1 | 8.98 | 0.003** |
| Secondary forest vs Fallow land | 1 | 8.17 | 0.008** |
| Primary forest vs Secondary forest | 1 | 0.86 | 0.652 |

\* $p \leq 0.05$ , \*\* $p \leq 0.01$ , \*\*\* $p \leq 0.001$

**Table S3:** Pairwise comparisons of dispersion (beta diversity) from centroid among habitat types based on the betadisper() analysis and Tukey’s HSD test.

| Pair comparisons | difference | lower | upper | p |
| --- | --- | --- | --- | --- |
| Primary forest vs Fallow land | -0.057 | -0.121 | 0.007 | 0.087 |
| Secondary forest vs Fallow land | -0.062 | -0.126 | 0.002 | 0.059 |
| Primary forest vs Secondary forest | -0.005 | -0.069 | 0.059 | 0.975 |

\* $p \leq 0.05$ , \*\* $p \leq 0.01$ , \*\*\* $p \leq 0.001$

**Table S4:** Pairwise Dunn’s test results for the comparison of the effect of habitat type on the normalized log ratio constant (lrc) from the rarefaction asymptotic curves.

| Pair comparisons | Z | p |
| --- | --- | --- |
| Primary forest - Secondary forest | 5.49 | < 0.001*** |
| Primary forest - Fallow land | -14.58 | < 0.001*** |
| Secondary forest - Fallow land | -9.11 | < 0.001*** |

\* $p \leq 0.05$ , \*\* $p \leq 0.01$ , \*\*\* $p \leq 0.001$

**Table S5:** Pairwise Dunn’s test results for the comparison of the effect of sampling method and habitat type on diversity indices. (a) Species richness (S), (b) Shannon index (H), and (c) Simpson index.

| Pair comparisons | Z | p |
| --- | --- | --- |
| Fallow land_Pitfall - Fallow land_Winkler | -0.36 | 0.832 |
| Fallow land_Pitfall - Primary forest_Pitfall | -1.00 | 0.528 |
| Fallow land_Winkler - Primary forest_Pitfall | -0.64 | 0.708 |
| Fallow land_Pitfall - Primary forest_Winkler | -3.58 | 0.005** |
| Fallow land_Winkler - Primary forest_Winkler | -3.22 | 0.006** |
| Primary forest_Pitfall - Primary forest_Winkler | -2.58 | 0.021* |
| Fallow land_Pitfall - Secondary forest_Pitfall | -0.12 | 0.902 |
| Fallow land_Winkler - Secondary forest_Pitfall | 0.23 | 0.874 |
| Primary forest_Pitfall - Secondary forest_Pitfall | 0.88 | 0.570 |
| Primary forest_Winkler - Secondary forest_Pitfall | 3.46 | 0.004** |
| Fallow land_Pitfall - Secondary forest_Winkler | -3.09 | 0.008** |
| Fallow land_Winkler - Secondary forest_Winkler | -2.73 | 0.016* |
| Primary forest_Pitfall - Secondary forest_Winkler | -2.09 | 0.069 |
| Primary forest_Winkler - Secondary forest_Winkler | 0.49 | 0.777 |
| Secondary forest_Pitfall - Secondary forest_Winkler | -2.96 | 0.009** |

\* $p \leq 0.05$ , \*\* $p \leq 0.01$ , \*\*\* $p \leq 0.001$

b) Shannon index
| Pair comparisons | Z | p |
| --- | --- | --- |
| Fallow land_Pitfall - Fallow land_Winkler | -0.33 | 0.857 |
| Fallow land_Pitfall - Primary forest_Pitfall | -1.01 | 0.518 |
| Fallow land_Winkler - Primary forest_Pitfall | -0.68 | 0.673 |
| Fallow land_Pitfall - Primary forest_Winkler | -3.59 | 0.005** |
| Fallow land_Winkler - Primary forest_Winkler | -3.26 | 0.006** |
| Primary forest_Pitfall - Primary forest_Winkler | -2.58 | 0.021* |
| Fallow land_Pitfall - Secondary forest_Pitfall | -0.19 | 0.908 |
| Fallow land_Winkler - Secondary forest_Pitfall | 0.14 | 0.891 |
| Primary forest_Pitfall - Secondary forest_Pitfall | 0.82 | 0.617 |
| Primary forest_Winkler - Secondary forest_Pitfall | 3.40 | 0.005** |
| Fallow land_Pitfall - Secondary forest_Winkler | -3.10 | 0.007** |
| Fallow land_Winkler - Secondary forest_Winkler | -2.77 | 0.014* |
| Primary forest_Pitfall - Secondary forest_Winkler | -2.08 | 0.070 |
| Primary forest_Winkler - Secondary forest_Winkler | 0.49 | 0.777 |
| Secondary forest_Pitfall - Secondary forest_Winkler | -2.90 | 0.011* |

\* $p \leq 0.05$ , \*\* $p \leq 0.01$ , \*\*\* $p \leq 0.001$

c) Simpson index
| Pair comparisons | Z | p |
| --- | --- | --- |
| Fallow land_Pitfall - Fallow land_Winkler | -0.33 | 0.857 |
| Fallow land_Pitfall - Primary forest_Pitfall | -1.01 | 0.518 |
| Fallow land_Winkler - Primary forest_Pitfall | -0.68 | 0.673 |
| Fallow land_Pitfall - Primary forest_Winkler | -3.59 | 0.005** |
| Fallow land_Winkler - Primary forest_Winkler | -3.26 | 0.006** |
| Primary forest_Pitfall - Primary forest_Winkler | -2.58 | 0.021* |
| Fallow land_Pitfall - Secondary forest_Pitfall | -0.19 | 0.908 |
| Fallow land_Winkler - Secondary forest_Pitfall | 0.14 | 0.891 |
| Primary forest_Pitfall - Secondary forest_Pitfall | 0.82 | 0.617 |
| Primary forest_Winkler - Secondary forest_Pitfall | 3.40 | 0.005** |
| Fallow land_Pitfall - Secondary forest_Winkler | -3.10 | 0.007** |
| Fallow land_Winkler - Secondary forest_Winkler | -2.77 | 0.014* |
| Primary forest_Pitfall - Secondary forest_Winkler | -2.08 | 0.070 |
| Primary forest_Winkler - Secondary forest_Winkler | 0.49 | 0.777 |
| Secondary forest_Pitfall - Secondary forest_Winkler | -2.90 | 0.011* |

**Table S6:** Permanova pairwise test results for the comparison of the effect of sampling method and habitat type on the community composition of ants.

| Pair comparisons | d.f. | F | p |
| --- | --- | --- | --- |
| Primary forest_Winkler vs Secondary forest_Winkler | 1 | 0.89 | 0.595 |
| Primary forest_Winkler vs Fallow land_Winkler | 1 | 5.60 | 0.005** |
| Primary forest_Winkler vs Secondary forest_Pitfall | 1 | 7.54 | 0.005** |
| Primary forest_Winkler vs Primary forest_Pitfall | 1 | 5.98 | 0.005** |
| Primary forest_Winkler vs Fallow land_Pitfall | 1 | 19.48 | 0.005** |
| Secondary forest_Winkler vs Fallow land_Winkler | 1 | 4.93 | 0.005** |
| Secondary forest_Winkler vs Secondary forest_Pitfall | 1 | 7.72 | 0.006** |
| Secondary forest_Winkler vs Primary forest_Pitfall | 1 | 6.41 | 0.006** |
| Secondary forest_Winkler vs Fallow land_Pitfall | 1 | 18.65 | 0.005** |
| Fallow land_Winkler vs Secondary forest_Pitfall | 1 | 4.28 | 0.005** |
| Fallow land_Winkler vs Primary forest_Pitfall | 1 | 3.93 | 0.005** |
| Fallow land_Winkler vs Fallow land_Pitfall | 1 | 4.41 | 0.005** |
| Secondary forest_Pitfall vs Primary forest_Pitfall | 1 | 0.88 | 0.595 |
| Secondary forest_Pitfall vs Fallow land_Pitfall | 1 | 5.89 | 0.005** |
| Primary forest_Pitfall vs Fallow land_Pitfall | 1 | 6.86 | 0.008** |

\* $p \leq 0.05$ , \*\* $p \leq 0.01$ , \*\*\* $p \leq 0.001$

**Table S7:** Pairwise comparisons of dispersion (beta diversity) from centroid among sampling methods and habitat types based on the betadisper() analysis and Tukey’s HSD test.

| Pair comparisons | difference | lower | upper | p |
| --- | --- | --- | --- | --- |
| Fallow land_Winkler-Fallow land_Pitfall | 0.177 | 0.080 | 0.273 | < 0.001*** |
| Primary forest_Pitfall-Fallow land_Pitfall | 0.162 | 0.066 | 0.259 | < 0.001*** |
| Primary forest_Winkler-Fallow land_Pitfall | -0.005 | -0.101 | 0.092 | 1 |
| Secondary forest_Pitfall-Fallow land_Pitfall | 0.163 | 0.066 | 0.259 | < 0.001*** |
| Secondary forest_Winkler-Fallow land_Pitfall | -0.010 | -0.107 | 0.087 | 1 |
| Primary forest_Pitfall-Fallow land_Winkler | -0.014 | -0.111 | 0.083 | 0.998 |
| Primary forest_Winkler-Fallow land_Winkler | -0.181 | -0.278 | -0.085 | < 0.001*** |
| Secondary forest_Pitfall-Fallow land_Winkler | -0.014 | -0.111 | 0.083 | 0.998 |
| Secondary forest_Winkler-Fallow land_Winkler | -0.187 | -0.283 | -0.090 | < 0.001*** |
| Primary forest_Winkler-Primary forest_Pitfall | -0.167 | -0.264 | -0.071 | < 0.001*** |
| Secondary forest_Pitfall-Primary forest_Pitfall | 0.000 | -0.097 | 0.097 | 1 |
| Secondary forest_Winkler-Primary forest_Pitfall | -0.173 | -0.269 | -0.076 | < 0.001*** |
| Secondary forest_Pitfall-Primary forest_Winkler | 0.167 | 0.071 | 0.264 | < 0.001*** |
| Secondary forest_Winkler-Primary forest_Winkler | -0.005 | -0.102 | 0.091 | 1 |
| Secondary forest_Winkler-Secondary forest_Pitfall | -0.173 | -0.269 | -0.076 | < 0.001*** |

\* $p \leq 0.05$ , \*\* $p \leq 0.01$ , \*\*\* $p \leq 0.001$

## Notes

https://doi.org/10.5281/zenodo.17809130

### Competing Interest Statement

The authors have declared no competing interest.

## REFERENCES

Agosti D & Alonso, L.E. 2000. The ALL protocol: A standard protocol for the collection of ground-dwelling ants. In: Agosti, D. et al. (Eds), Ants: Standard methods for measuring and monitoring biodiversity: 280. Smithsonian Institution Press.

Agosti, D. & Alonso, L.E. 2003. El protocolo ALL: Un estándar para la colección de hormigas del suelo. In: F.F. (Ed.), Introducción a las hormigas de la región Neotropical: 424.

Alongo, L.S. 2007. Etude de l’effet des lisières sur l’humidité équivalente et la température du sol d’un écosystème forestier de la cuvette centrale congolaise. Mémoire DEA, Université de Kisangani, République Démocratique du Congo.

Alongo, S. 2013. Etude microclimatique et pédologique de l’effet de lisière en Cuvette centrale congolaise: impact écologique de la fragmentation des écosystèmes — cas des séries Yangambi et Yakonde à la région de Yangambi (R.D. Congo). Thèse, Université Libre de Bruxelles.

Andersen, A.N., Hoffmann, B.D., Müller, W.J. & Griffiths, A.D. 2002. Using ants as bioindicators in land management. Journal of Applied Ecology 39: 8–17. 10.1046/j.1365-2664.2002.00704.x

Ambakina, H. 2022. Diversité entomologique et arborescente dans les forêts du village Bangbasende (Tshopo, RD Congo). Mémoire DES, Université de Kisangani.

Andersen, A.N. & Majer, J.D. 2004. Ants show the way down under: invertebrates as bioindicators in land management. Frontiers in Ecology and the Environment 2: 291–298.

Andersen, A.N., van Ingen, L.T. & Campos, R.I. 2007. Contrasting rainforest and savanna ant faunas in monsoonal northern Australia: a rainforest patch in a tropical savanna landscape. Australian Journal of Zoology 55: 363–369.

Bastos, A.H.S. & Harada, A.Y. 2011. Leaf-litter amount as a factor in the structure of a ponerine ant community in an eastern Amazonian rainforest, Brazil. Revista Brasileira de Entomologia 55: 589–596.

Beck, H.E. et al. 2018. Present and future Köppen–Geiger climate classification maps at 1-km resolution. Scientific Data 5: 180214. 10.1038/sdata.2018.214

Benson, W. & Harada, A.Y. 1988. Local diversity of tropical and temperate ant faunas. Acta Amazonica 18: 275–289.

Blaimer, B.B., Brady, S.G., Schultz, T.R., Lloyd, M.W., Fisher, B.L. & Ward, P.S. 2015. Phylogenomic methods outperform traditional multi-locus approaches in resolving deep evolutionary history: a case study of formicine ants. BMC Evolutionary Biology 15: 271. 10.1186/s12862-015-0552-5

Bolton, B. 2000. The ant tribe Dacetini (Hymenoptera, Formicidae). Memoirs of the American Entomological Institute 65: 1–1028.

Boyle, M.J.W. et al. 2025. Causes and consequences of insect decline in tropical forests. Nature Reviews Biodiversity 1: 315–331.

Bronstein, J.L., Alarcón, R. & Geber, M. 2006. The evolution of plant–insect mutualisms. New Phytologist 172: 412–428.

Daily, G.C., Ehrlich, P.R. & Sanchez-Azofeifa, G.A. 2001. Countryside biogeography: use of human-dominated habitats by the avifauna of southern Costa Rica. Ecological Applications 11: 1–13.

Dejean, A., Corbara, B., Orivel, J. & Leponce, M. 2007. Rainforest canopy ants: the implications of territoriality and predatory behavior. Functional Ecosystems and Communities 1: 105–120.

Delabie, J.H.C. & Fowler, H.G. 1995. Soil and litter cryptic ant assemblages in Bahian cocoa plantations. Pedobiologia 39: 423–433.

Devineau, J.-L. 1975. Étude quantitative des forêts galeries de Lamto (Moyenne Côte d’Ivoire). Thèse de 3e cycle, Université Paris 6.

Eguchi, K., Bui, T.V., Yamane, S. & Terayama, M. 2015. Redefinition of the genus Bannapone and description of B. cryptica sp. nov. (Hymenoptera: Formicidae: Amblyoponinae). Zootaxa 4013: 77–86.

Emery, C. 1911. Hymenoptera. Fam. Formicidae. Subfam. Ponerinae. Genera Insectorum 118: 1–125.

FAO. 2010. Global Forest Resources Assessment 2010. Food and Agriculture Organization of the United Nations, Rome.

Fimbel, R.A., Grajal, A. & Robinson, J.G. 2001. Logging-wildlife issues in the tropics: an overview. In: Fimbel, R.A., Grajal, A. & Robinson, J.G. (Eds), The Cutting Edge: Conserving Wildlife in Logged Tropical Forests. Columbia University Press.

Fisher, B.L. & Bolton, B. 2016. Ants of Africa and Madagascar: A guide to the genera (Hymenoptera, Formicidae). University of California Press.

Giam, X. 2017. Global biodiversity loss from tropical deforestation. PNAS 114(23): 5775–5777.

Gotelli, N.J. & Colwell, R.K. 2001. Quantifying biodiversity: procedures and pitfalls in the measurement and comparison of species richness. Ecology Letters 4: 379–391.

Groc, S. 2007. Structure taxonomique et application de la notion de minimalisme taxonomique à la myrmécofaune des Nouragues, Guyane française. Mémoire Master 2 SEP, Université Paris 6 & MNHN.

Groc, S., Orivel, J., Dejean, A., Martin, J.M., Etienne, M.P., Corbara, B. & Delabie, J.H.C. 2009. Baseline study of the leaf-litter ant fauna in a French Guianese forest. Insect Conservation and Diversity 2: 183–193.

Hölldobler, B. & Wilson, E.O. 1990. The Ants. Harvard University Press.

Karsenty, A. 2012. Le rôle de l’agriculture dans la déforestation et la dégradation en RDC: situation actuelle, perspectives et solutions possibles. Online document.

Kaspari, M., O’Donnell, S. & Kercher, J.R. 2000. Energy, density, and constraints to species richness: ant assemblages along a productivity gradient. American Naturalist 155: 280–293.

Kouakou, L.M.M. 2009. Impact des activités humaines sur la diversité biologique des fourmis terricoles du parc national du Banco. Mémoire de maîtrise, Université d’Abobo-Adjamé.

LaPolla, J.S., Suman, T., Sosa-Calvo, J. & Schultz, T.R. 2006. Leaf-litter ant diversity in Guiana. Biodiversity and Conservation 16: 491–510.

Laso Bayas, J.C., et al. 2022. Drivers of tropical forest loss between 2008 and 2019. Scientific Data 9: 146.

Lassau, S.A. & Hochuli, D.F. 2004. Effects of habitat complexity on ant assemblages. Ecography 27: 157–164.

Leponce, M., Delabie, J.H.C., Orivel, J., Jacquemin, J., Calvo Martín, M. & Dejean, A. 2019. Tree-dwelling ant survey in Mitaraka, French Guiana. In: Touroult, J. (Ed.), Our Planet Reviewed: 2015 large-scale biotic survey in Mitaraka, French Guiana. Zoosystema 41: 163–179. 10.5252/zoosystema2019v41a10

Lévieux, J. 1983. Soil fauna of tropical savannahs: 4. The ants. In: Bourlière, F. (Ed.), Tropical Savannahs: 525–540. Elsevier, Amsterdam.

Li, X. 2025. Soil microbial communities in tropical rainforests: diversity, functionality, and environmental implications. Journal of Forest Research 14: 557.

Majer, J.D., Orabi, G. & Bisevac, L. 2007. Ants (Hymenoptera: Formicidae) pass the bioindicator scorecard. Myrmecological News 10: 69–76.

Marsh, C.J. et al. 2025. Tropical forest clearance impacts biodiversity and function, whereas logging changes structure. Science 387(6730): 171–175.

Martelli, M.G., Ward, M.M. & Fraser, A.M. 2004. Ant diversity sampling on the Southern Cumberland Plateau: a comparison of litter sifting and pitfall trapping. Southeastern Naturalist 3(1): 113–126.

Mayaux, P. et al. 2003. Évolution du couvert forestier du bassin du Congo mesurée par télédétection spatiale. Bois et Forêts des Tropiques 277: 45–52.

Mayer, J.D. & Delabie, J.H.C. 1994. Comparison of the ant communities of annually inundated and terra firme forests at Trombetas in the Brazilian Amazon. Insectes Sociaux 41: 343–359.

McCoy, E.D. & Bell, S.S. 1991. Habitat structure: the evolution and diversification of a complex topic. In: McCoy, E.D., Bell, S.S. & Mushinsky, H.R. (Eds), Habitat structure: The physical arrangements of objects in space: 3–27. Chapman & Hall.

Mertl, A.L., Ryder Wilkie, K.T.R. & Traniello, J.F.A. 2009. Impact of flooding on the species richness, density and composition of Amazonian litter-nesting ants. Biotropica 41: 633–641.

Pacheco, R. & Vasconcelos, H.L. 2012a. Habitat diversity enhances ant diversity in a naturally heterogeneous Brazilian landscape. Biodiversity and Conservation 21: 797–809. 10.1007/s10531-011-0221-y

Pacheco, R. & Vasconcelos, H.L. 2012b. Subterranean pitfall traps: is it worth including them in your ant sampling protocol? Psyche 2012: 870794. 10.1155/2012/870794

Philpott, S.M. & Armbrecht, I. 2006. Biodiversity in tropical agroforests and the ecological role of ants and ant diversity in predatory function. Ecological Entomology 31: 369–377.

Pillay, R., Venter, M., Aragon-Osejo, J., González-del-Pliego, P., Hansen, A.J., Watson, J.E.M. & Venter, O. 2022. Tropical forests are home to over half of the world’s vertebrate species. Frontiers in Ecology and the Environment 20(1): 10–15. 10.1002/fee.2420

Rohr, J.R., Mahan, C.G. & Kim, K.C. 2006. Developing a monitoring program for invertebrates: guidelines and a case study. Conservation Biology 21: 422–433.

Schultheiss, P., Nooten, S.S., Wang, R., Wong, M.K.L., Brassard, F. & Guénard, B. 2022. The abundance, biomass, and distribution of ants on Earth. PNAS 119(40): e2201550119. 10.1073/pnas.2201550119

Silva, F.H.O., Delabie, J.H.C., dos Santos, G.B., Meurer, E. & Marques, M.I. 2013. Mini-Winkler extractor and pitfall traps as complementary methods to sample Formicidae. Neotropical Entomology 42: 351–358.

Stein, A., Gerstner, K. & Kreft, H. 2014. Environmental heterogeneity as a universal driver of species richness across taxa, biomes and spatial scales. Ecology Letters 17: 866–880. 10.1111/ele.12277

Stein, A. & Kreft, H. 2015. Terminology and quantification of environmental heterogeneity in species-richness research. Biological Reviews 90: 815–836. 10.1111/brv.12135

Strona, G. & Bradshaw, C.J.A. 2022. Coexistence dominates future vertebrate losses from climate and land use change. Science Advances 8(50).

Tilman, D., Clark, M., Williams, D.R., Kimmel, K., Polansky, S. & Packer, C. 2017. Future threats to biodiversity and pathways to their prevention. Nature 546: 73–81.

Underwood, E.C. & Fisher, B.L. 2006. The role of ants in conservation monitoring: if, when, and how. Biological Conservation 132: 166–182.

Vasconcelos, H.L. & Vilhena, J.M.S. 2006. Species turnover and vertical partitioning of ant assemblages in the Brazilian Amazon. Biotropica 38: 100–106. 10.1111/j.1744-7429.2006.00113.x

Ward, P.S. & Fisher, B.L. 2016. Tales of Dracula ants: the evolutionary history of the ant subfamily Amblyoponinae (Hymenoptera: Formicidae). Systematic Entomology 41: 683–693. 10.1111/syen.12186

WWF. 2007. Climate change impacts in the Amazon: review of scientific literature. WWF report.

Yangambi Research Station. n.d. Yangambi.org.

Yeo, K., Delsinne, T., Konaté, S., Alonso, L.L., Aïdara, D. & Peeters, C. 2017. Diversity and distribution of ant assemblages above and below ground in a West African forest–savannah mosaic (Lamto, Côte d’Ivoire). Insectes Sociaux 64: 155–168. 10.1007/s00040-016-0527-6

York, A. 2000. Long-term effects of frequent low-intensity burning on ant communities in coastal blackbutt forests of southeastern Australia. Australian Ecology 25: 83–98.

Yoshimura, M. & Fisher, B.L. 2012. A revision of male ants of the Malagasy Amblyoponinae (Hymenoptera: Formicidae) with resurrections of the genera Stigmatomma and Xymmer. PLoS ONE 7: e33325. 10.1371/journal.pone.0033325

